# Multi-drug resistant *E. coli* displace commensal *E. coli* from the intestinal tract, a trait associated with elevated levels of genetic diversity in carbohydrate metabolism genes

**DOI:** 10.1101/2022.11.25.517930

**Authors:** Christopher H. Connor, Amanda Z. Zucoloto, Ian-Ling Yu, Jukka Corander, Braedon McDonald, Alan McNally

**Author notes:** Corresponding authors: Prof Alan McNally, Institute of Microbiology and Infection, College of Medical and Dental Science, University of Birmingham, Edgbaston, Birmingham, United Kingdom, B15 2TT. and Dr Braedon McDonald, Department of Critical Care Medicine, University of Calgary, Calgary, Alberta, Canada.

## Abstract

Extra-intestinal pathogenic *E. coli* (ExPEC) can cause a variety of infections outside of the intestine and are a major causative agent of urinary tract infections. Treatment of these infections is increasingly frustrated by antimicrobial resistance (AMR) diminishing the number of effective therapies available to clinicians. Incidence of multi-drug resistance (MDR) is not uniform across the phylogenetic spectrum of *E. coli*. Instead AMR is concentrated in select lineages, such as ST131, which are MDR pandemic clones that have spread AMR globally. Using a gnotobiotic mouse model we demonstrate that an MDR *E. coli* ST131 is capable out-competing and displacing non-MDR *E. coli* from the gut *in vivo*. This is achieved in the absence of antibiotic treatment mediating a selective advantage. In mice colonised with non-MDR *E. coli* strains, challenge with MDR *E. coli* either by oral gavage or co-housing with MDR *E. coli* colonized mice results in displacement and dominant intestinal colonization by MDR *E. coli* ST131. To investigate the genetic basis of this superior gut colonization ability by MDR *E. coli*, we used a functional pangenomic analysis of 19,571 *E. coli* genomes revealing that carriage of AMR genes is associated with increased diversity in carbohydrate metabolism genes. The data presented here demonstrate that independent of antibiotic selective pressures, MDR *E. coli* display a competitive advantage to colonise the mammalian gut and points to a vital role of metabolism in the evolution and success of MDR lineages of *E. coli* via carriage and spread.

## Introduction

Infections by multi-drug resistant (MDR) Gram-negative pathogens now represent one of the greatest global public health challenges of our generation, with the World Health Organisation declaring them of utmost international importance. Chief amongst these pathogens is MDR *Escherichia coli*, which are responsible for an alarming rise in the incidence of antimicrobial resistant (AMR) blood stream and urinary tract infections^1^. MDR in *E. coli* is heavily associated with the carriage of large MDR plasmids carrying genes encoding extended spectrum beta-lactamases (ESBL) and carbapenemases such as NDM, KPC, and Oxa-48 which confer resistance to 3^rd^ generation cephalosporins and carbapenem classes of antibiotics respectively^1^. Intriguingly such plasmids are very rarely found in intestinal pathogenic *E. coli* such as *E. coli* O157 or common enteropathogenic and enterotoxigenic *E. coli* strains. Rather MDR plasmid carriage is concentrated in a number of lineages responsible for extra-intestinal pathogenesis such as blood stream and urinary tract infections^2^.

Extra-intestinal pathogenic *E. coli* (ExPEC) is the name given to *E. coli* strains capable of causing extra-intestinal infections, but do not represent a phylogenetically distinct group of organisms. Rather ExPEC are found across the species phylogeny mainly in phylogroups B2, D, and F^3^. Recent longitudinal surveys of national blood stream infection isolates have shown the most common ExPEC lineages to be ST131, ST73, ST69, ST95, ST410, and members of the ST10 complex including ST167^4,5^. Equally as intriguingly, MDR plasmids are not evenly distributed amongst these ExPEC lineages but rather their carriage is concentrated in a small number of highly successful, globally disseminated clones^2^. The most successful of these clones is clade C of ST131, the most common cause of MDR blood stream and urinary tract infections worldwide^6^. Other common MDR *E. coli* lineages include ST69, ST410, ST167, and ST648^2^.

What makes certain clones or lineages of *E. coli* successful MDR pathogens is an ongoing question. Recent analysis of longitudinal blood stream infection isolates from Norway shows that ST131 strains successfully emerged to be dominant in the absence of MDR plasmid carriage indicating MDR alone is not the driver of their success^5^. Analysis of longitudinal UK isolates shows that MDR alone is not sufficient to drive strains to complete dominance of the epidemiological landscape^4^. Recent evidence suggests an important phenotype that differentiates MDR *E. coli* from other lineages is their ability to rapidly and asymptomatically colonise the intestinal tract of humans. The COMBAT study of 1847 people travelling from the Netherlands to Asia, Africa and South America found that 34% of those travelling acquired an ESBL *E. coli* in their intestinal tract during their journey, with that number increasing to 75% of those travelling to Asia^7^. Of those colonised in the study, 11% were colonised for up to 12 months. A small scale study of University of Birmingham students found that all participants travelling to Asia became colonised by an ESBL *E. coli*, with genomic analysis confirming that this was due to acquisition of a new MDR strain and not the resident commensal *E. coli* becoming MDR^8^. A recent study of medical personnel travelling to Laos sampled travellers in real time to show that every single person was colonised by an MDR *E. coli* during travel, with colonisation occurring within days after arrival^9^. A very recent study by our group deploying metagenomic sequencing on the COMBAT study samples showed that when people were colonised by an MDR *E. coli* there was no detectable impact on diversity or composition of the wider gut microbiome as a result of MDR colonisation^10^. Whilst observational studies have yielded hypothesis-generating data suggesting stain-to-strain competition, this has never been directly tested in vivo.

Studies investigating genetic determinants differentiating MDR *E. coli* lineages from the rest of the population have also uncovered a number of parallel observations. Studies comparing clade C ST131 to its ancestral population show a collection of adaptive nucleotide substitutions in chromosomal promoter regions associated with the specific plasmid caried by the strain^11^, a pattern which has also been seen in ST167^12^. A high resolution study of ST131 using a pangenome approach to identify allelic variations in genes found a highly elevated number of nucleotide substitutions in genes involved in mammalian colonisation including anaerobic metabolism, iron acquisition, and adhesins. This pattern was not seen in successful non-MDR ExPEC lineages such as ST73^13^. Such allelic diversity in anaerobic metabolism genes has also been seen in ST167 and ST648^12,14^ and it was also shown that recombination of new alleles of *fhu* iron acquisition genes was the key evolutionary event underpinning the emergence the carbapenem resistant B4/H24RxC clone in the ST410 lineage^15^. We hypothesise that these genetic adaptations may contribute to more effective colonisation of the mammalian gut.

Here we use a gnotobiotic mouse model of intestinal colonisation to directly test the hypothesis that MDR *E. coli* can outcompete commensal *E. coli* to establish dominant colonization the intestinal tract *in vivo* in the absence of any antibiotic selection. We demonstrate that an MDR ST131 strain can out-compete a commensal strain to establish intestinal colonization in gnotobiotic mice.

Furthermore, when introduced into the gut of mice pre-colonized with commensal *E. coli*, MDR ST131 could displace commensal *E. coli* from the gut to establish dominant colonization. Displacement of the resident commensal strain occurs within 48 hours of cohousing with mice colonized by MDR ST131, with ST131 becoming the dominant strain in all co-housed mice. We also show that colonization with MDR *E. coli* ST131 stimulated a distinct pro-inflammatory cytokine response in the host caecum compared to mice colonized by a commensal strain. We then utilised a curated dataset of 19,571 genomes representing the full phylogenetic and AMR diversity of *E. coli* to identify a significant link between metabolism and carriage of AMR genes. We find that MDR lineages of *E. coli* display an increased nucleotide diversity in genes associated with core energy metabolism and carbohydrate utilisation which may afford competitive advantage for colonizing the mammalian gut compared to commensal *E. coli* strains.

## Methods

### Mouse colonisation

Mouse colonisation experiments were conducted at the International Microbiome Centre, University of Calgary, Canada. Germ-free mice were bred and maintained in flexible film isolators in our axenic breeding facility, and germ-free status was confirmed by a combination of Gram staining, Sytox green DNA staining, anaerobic and aerobic culture, and 16s rRNA gene amplicon sequencing from faeces. were maintained in isocages in our gnotobiotic facility. Germ-free C57BL/6 mice were colonised with 10^9^ CFU of bacteria via oral gavage (see table S1 and Fig S1&2 for details of strains used for colonisation) and maintained in sterile isocages in our gnotobiotic facility throughout the duration of experiments. Bacterial colonisation was monitored by CFU enumeration on UTI Chromogenic Agar (Thermo) from faecal pellets as well as by DNA extraction and strain specific probe-based qPCR. DNA from faecal pellets was extracted using the MagMAX Microbiome Ultra Nucleic Acid isolation kit (Thermo) on a KingFisher Flex instrument (Thermo) following manufacturer’s instructions. Strain specific primers and probes were designed to target unique genes (commensal: *clpP*, ST73: *lon*, ST131: *prtR* – Table S2) identified from genome data. Minimal cross reaction was observed in samples from monocolonised mice (Fig S3). Probes were manufactured by Integrated DNA Technologies (IDT). Reactions were performed using PrimeTime Gene Expression master mix (IDT) on a QuantStudio 1 system (Thermo). Reactions were performed following manufacturer’s recommended parameters: 3 minutes at 95*C, 40 cycles of 15 seconds at 95*C, 1 minute at 60*C, fluorescence readings taken at the end of the extension stage. A standard curve was generated from a bacterial culture of known CFU (Fig S4).

### Cytokine expression, histology and *in vitro* supernatant cultures

Sections of mouse gut were preserved in RNAlater Stabilization solution (Thermo) before homogenization in lysis buffer. Tissues were homogenised in RLT buffer (Qiagen) using a stainless-steel bead and agitating in a FastPrep-24 (MPBio) in 3×1 minute bursts at a speed setting of 6.0m/s with 1 minute intervals on ice. The resulting lysate was used as input for the RNeasy Mini kit (Qiagen) including an on-column DNase treatment (Qiagen). Extracted RNA was converted to cDNA using High-Capacity cDNA Reverse Transcription kit (Applied Biosystems) with random primers. Pre-designed probes for qPCR were obtained from Integrated DNA Technologies (IDT – Table S2) and reactions were performed using PrimeTime Gene Expression master mix (IDT) on a QuantStudio 1 system (Thermo). Reactions were performed following manufacturer’s recommended parameters: 3 minutes at 95*C, 40 cycles of 15 seconds at 95*C, 1 minute at 60*C, fluorescence readings taken at the end of the extension stage. Cytokine Ct values were normalised to the endogenous Pol2ra control and presented as deltaCt values. Significance was determined using 2-way ANOVA with Tukey’s multiple correction.

Sections of mouse gut were ‘swiss-rolled’, 10 % formalin fixed, dehyrated and embedded in paraffin. Tissues were sectioned longitudinally, mounted onto glass slides and stained with haematoxylin and eosin. Slides were imaged using an Axioscan 7 Slide Scanner (Zeiss) and blind scored using Fiji following the scoring system in^16^.

Bacteria were grown in LB broth to an OD600 of between 0.4 and 0.6, cultures were pelleted at 4,000rpm for 15 minutes. The supernatant was passed through a 0.22μm filter, the filtrate was diluted in a 1 to 1 ratio with fresh LB. Bacteria were inoculated into the supernatant mixture and grown in a 96 well plate at 37°C with growth measured by OD600 readings at 10 minute intervals by a Spark Microplate Reader (Tecan)

### Genomic dataset curation, pangenome construction and AMR gene detection

A total of 20 lineages, or sequence types (ST) were selected from the literature with a focus on ExPEC lineages but also including EHEC and EPEC clones. This resulted in a dataset of 19,571 *E. coli* genome sequences encompassing all the *E. coli* phylogroups (A, B1, B2, D, E and F/G) (Table S3) [10.6084/m9.figshare.c.6147189]. The earliest samples with reliable metadata were from 1980 however the majority were sequenced in recent decades [Fig S5]. Humans represented the major source niche for all lineages except ST117 for which poultry was the major niche (Fig S5). This dataset contained samples from multiple countries however Europe and North America accounted for the majority (Fig S6).

Genome assemblies for each lineage were downloaded from Enterobase^17^ using a custom python script [https://github.com/C-Connor/EnterobaseGenomeAssemblyDownload]. Duplicated assemblies were identified using Mash v1.1.1^18^ to estimate genome similarity, a custom R script then removed isolates with a Mash distance of 0 [https://github.com/C-Connor/MashDistDeReplication]. Dendrograms of Mash distances were also constructed and examined for outlier genomes that were not part of a larger cluster. The remaining genome files were annotated with Prokka v1.12^19^ and pangenomes were constructed with Roary v3.10.2^20^ using a 95% identity threshold, a 99% core genome threshold, paralog splitting was disabled, and a core genome alignment was produced using MAFFT. AMR genes were detected using Abricate v 0.8 [https://github.com/tseemann/abricate] with the Resfinder-2018 database^21^, results were filtered to remove hits with less than 80% resistance gene coverage. A phylogeny of the whole dataset was constructed using MashTree v0.36.2^22^ and visualised in iTOL^23^.

### Pangenome functional annotation

Pangenome reference Fasta files produced by Roary were functionally annotated using emapper-1.0.3-3-g3e22728^24^ based on eggNOG orthology data^25^. Sequence searches were performed using DIAMOND^26^. Functional annotation data was combined with the gene presence absence matrix and analysed in R v4.0.3. To examine if there was any association between functional composition of the accessory pangenome or core genome linear regression was performed between carriage of AMR (as a proportion of the population with 2 or more AMR genes) and individual COG categories, correcting for multiple testing.

## Results

### Non-MDR ExPEC and MDR ExPEC are efficient colonisers of germ-free mice, with both ExPEC outcompeting a commensal isolate in co-colonisation

To determine whether MDR *E. coli* is able to out-compete non-MDR *E. coli* in the intestine *in vivo*, we performed competitive colonization experiments in GF mice using three different strains of *E. coli:* a non-MDR potential pathogen (ST73) commensal strain isolated from a healthy human volunteer; a pathogenic non-MDR ExPEC (ST73) strain isolated from a bacteraemia patient; and a pathogenic MDR ExPEC (ST131) strain isolated from a bacteraemia patient (Table S1). All three strains possessed an equivalent virulence associated gene profile, however the MDR ExPEC strain possessed a greater number of AMR genes (Fig S1 and S2). Mice were inoculated with 10^9^ CFU of each strain via oral gavage and bacterial growth was measured by enumeration of CFU from the faeces as well as strain specific qPCR. Under these conditions all strains could individually monocolonise germ free mice (Fig S3). Competitive colonization between strains was investigated by orally gavaging GF mice with a 1:1 ratio of 2 strains in combination (see Fig 1A for combinations) followed by quantification of each strain in the faeces over the subsequent week. Both the non-MDR and MDR ExPEC strains out-competed the commensal achieving 96.1% and 80.7% of the total growth respectively by day 6 post gavage (Fig 1A). Neither the non-MDR ExPEC nor the MDR ExPEC was able to out-compete the other with MDR ExPEC accounting for 47.1% of the growth by day 6 (Fig 1A).

**Figure 1.**
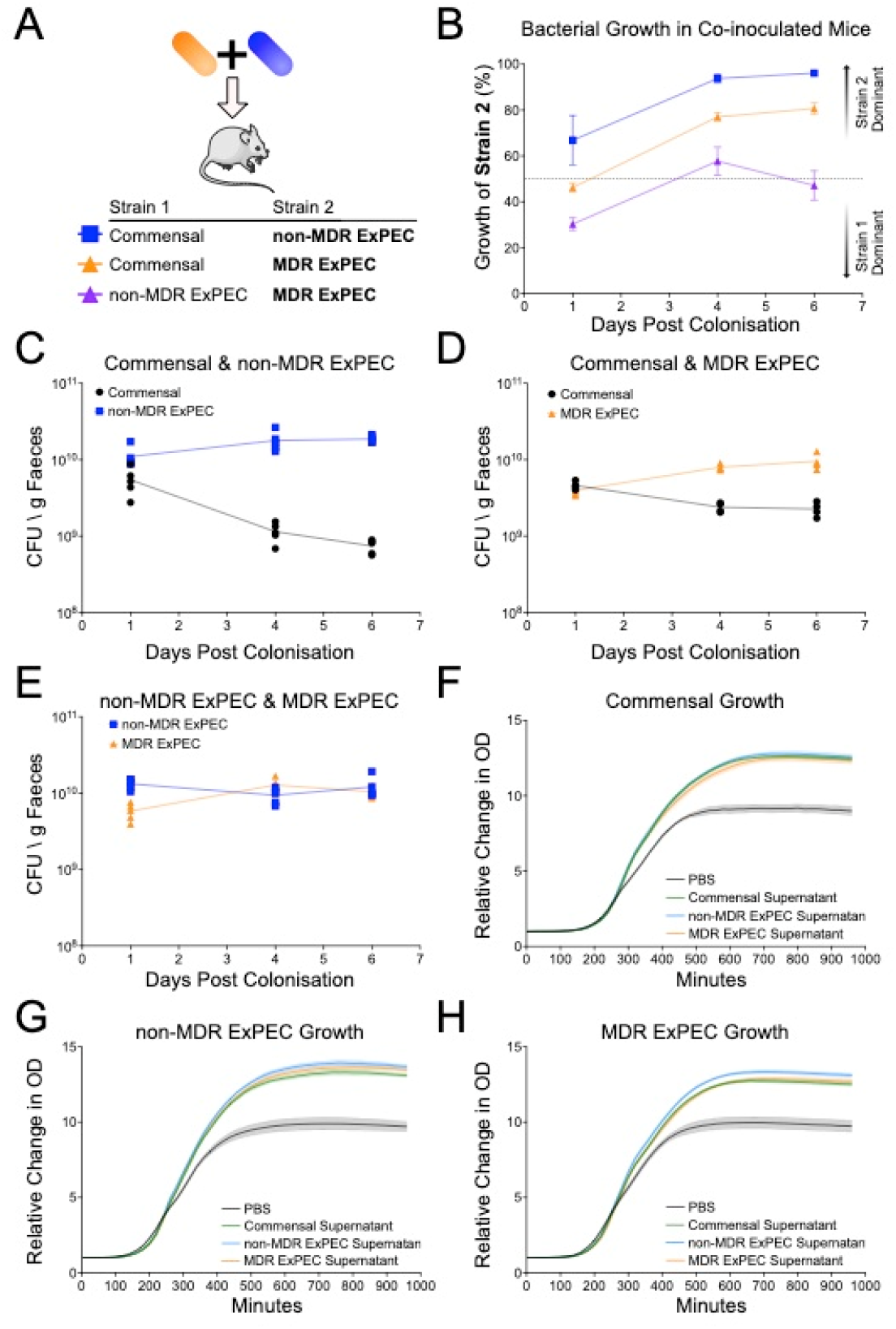
Co-colonisation of mice with 2 strains of *E. coli* and supernatant growth curves. A, Schematic of mouse colonisation set up, two strains of *E. coli* are mixed immediately prior to gavage. B, Growth of *E. coli* in co-colonised mice measured by strain specific qPCR. Growth of the Strain 2 listed in the schematic is plotted as a percentage of total growth against days post gavage (n=5, commensal & non-MDR ExPEC, growth of non-MDR ExPEC in blue; n=4, commensal & MDR ExPEC, growth of MDR ExPEC in orange, n=5 non-MDR ExPEC & MDR ExPEC, growth of MDR ExPEC in purple). Dotted line at 50% indicating equivalent growth of both strains, values higher than 50% indicate higher growth of strain 2 while values less than 50% indicate lower growth of strain 2. C, D, E, CFU per gram of faeces as measured by qPCR for mice colonised simultaneously by a commensal (black) & non-MDR ExPEC (blue) strain (C), commensal (black) & MDR ExPEC (orange) (D) or non-MDR ExPEC (blue) & MDR ExPEC (orange) (E). F, G, H, Growth of the commensal (F), non-MDR ExPEC (G) or MDR ExPEC (H) strain in 50% LB + 50% PBS (black), commensal supernatant (green), non-MDR ExPEC supernatant (blue) or MDR ExPEC supernatant (orange), (n=6).

It is possible that the non-MDR ExPEC and MDR ExPEC are out-competing the commensal strain due to phage or production of a secreted toxin. To test for this we grew each strain in LB broth overnight, filter sterilised the spent medium before mixing in a 1:1 ratio with fresh LB, and investigated the growth kinetics of all strains in the presence of supernatant from competing strains (Fig. 1 F-H). No strain displayed any impairment in growth in the presence of either autologous or heterologous supernatant indicating that no strains were releasing toxins / phage to kill competing strain (Fig. 1F-H).

### MDR ExPEC efficiently displaces an established commensal from the mouse intestinal tract

Next, we aimed to determine whether MDR *E. coli* could displace commensal *E. coli* that had already established a colonization niche within the intestine. Germ-free mice were colonised with a commensal strain by oral gavage and colonisation allowed to stabilise for one week before challenging with a second *E. coli* strain (Fig 2A). When commensal-colonized mice were challenged with MDR ExPEC strain, it rapidly out-competes the resident commensal strain within 4 days accounting for greater than 60% of the CFU (Fig 2B and C). By day 21 the MDR ExPEC strain accounted for 80.4% of the CFU in the faeces. In contrast, this displacement is not observed when commensal-mice colonised are challenged with the non-MDR ExPEC strain, which results in an equilibrium of 50-50 colonisation of both strains within 4 days and remains equivalently co-colonized at 21 days post-challenge (Fig 2B and D). Conversely, when MDR ExPEC monocolonized mice are challenged with a commensal strain, the commensal is unable to displace the resident MDR ExPEC strain, accounting for 20.4% of the CFU at day 4 and further diminishing to 15.3% at day 21 (Fig 2B and E). These results are not due to changes in commensal colonisation as growth remains stable in the control group throughout the experiment (Fig 2F). Collectively, these data demonstrate that MDR *E. coli* is able to displace resident commensal *E. coli* to established dominant colonization of the gut *in vivo*.

**Figure 2.**
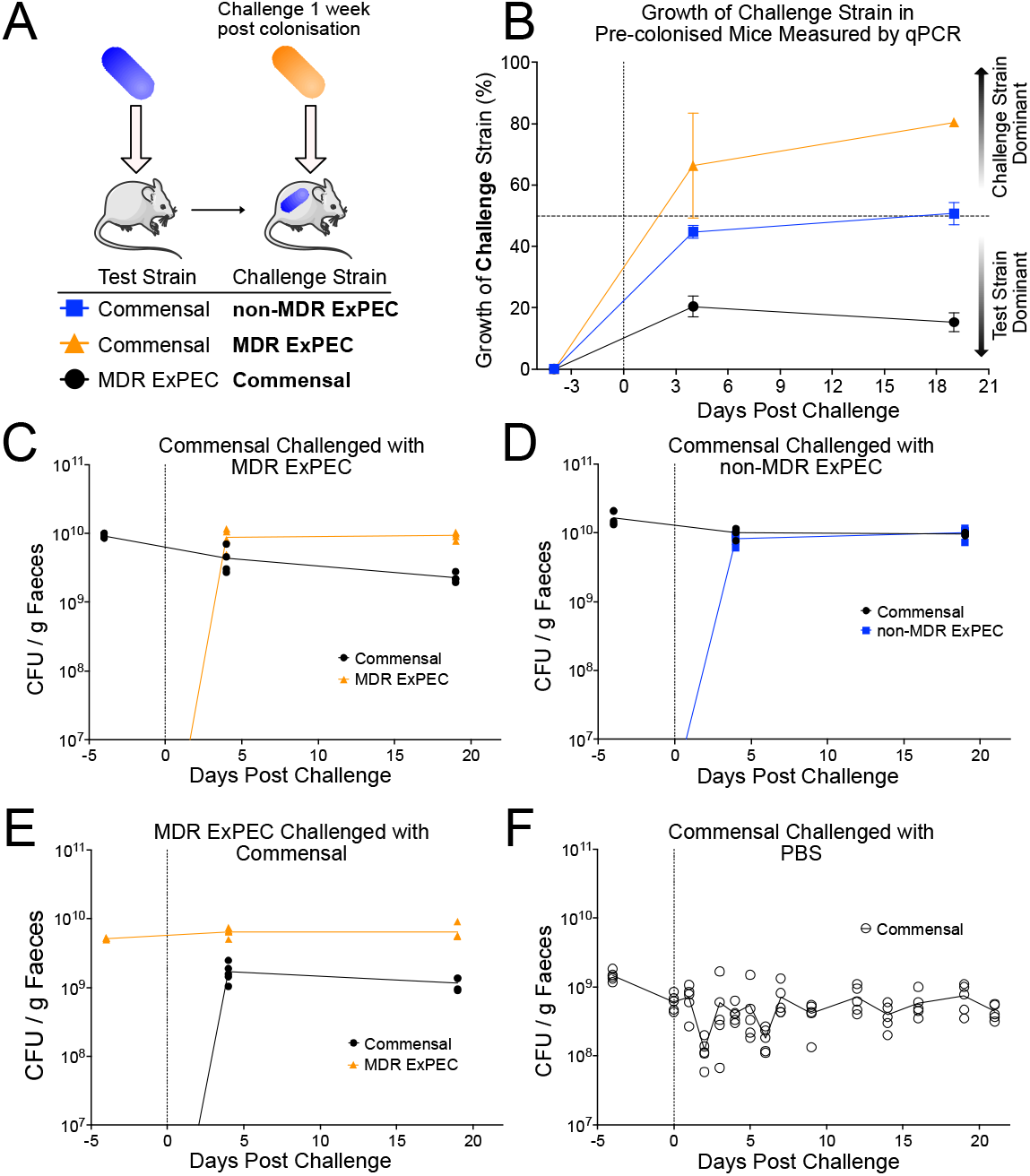
Mice colonised with a test strain for 1 week before challenge with a second strain. A, Schematic of displacement experiment in which mice are pre-colonised with a 10^9^ CFU of the test strain followed a week later by a second gavage with 10^9^ CFU of a challenge strain. B, Growth of the challenge strain described in panel A in pre-colonised mice measured by strain specific qPCR at select timepoints. Data presented as percentage of total growth attributable to challenge strain (n=5 for commensal vs. non-MDR ExPEC, growth of non-MDR ExPEC in blue, n=4 for commensal vs. MDR ExPEC, growth of MDR ExPEC in orange, n=4 for MDR ExPEC vs. commensal, growth of commensal in black). Vertical dotted line indicates timepoint for challenge with second strain. Horizontal dashed line indicates 50% corresponding to equivalent colonisation of test and challenge strains. C-E, Growth of both colonising strains in mice measured by strain specific qPCR, CFU calculated against a standard curve and normalised to faecal pellet weight. Growth of commensal strain in black circles, growth of non-MDR ExPEC in blue squares and growth of MDR ExPEC in orange triangles. Vertical dotted line indicates time of challenge with second strain. F, Growth of commensal in colonised mice challenged with PBS. Growth measured by CFU enumeration from plates at regular intervals. Vertical dotted line indicates time of challenge with PBS.

To determine whether our findings using oral gavage (with large quantities of bacteria) were replicated in the setting of environmental exposure to MDR *E. coli*, we co-housed commensal *E. coli* monocolonized mice and MDR ExPEC monocolonized mice (Fig 3A). Within 1 week of co-housing, MDR ExPEC is observed in the faeces of all mice, and in fact becomes the dominant coloniser in all mice (Fig 3). By day 4 of co-housing, MDR ExPEC accounts for nearly 80% of the CFU in all co-housed mice, and this colonization dominance is persistent and sustained for many weeks (Fig. 3B). Of note, MDR ExPEC - monocolonized mice did acquire a low level of colonization by commensal *E. coli* following co-housing, but MDR ExPEC remained the dominant strain with commensal accounting for less than 20% of faecal CFUs. This MDR ExPEC phenotype is not due to strain specific differences in inherent growth rate, as the total CFU recovered from MDR ExPEC monocolonised mice is equivalent to commensally monocolonised mice (Fig S3).

**Figure 3.**
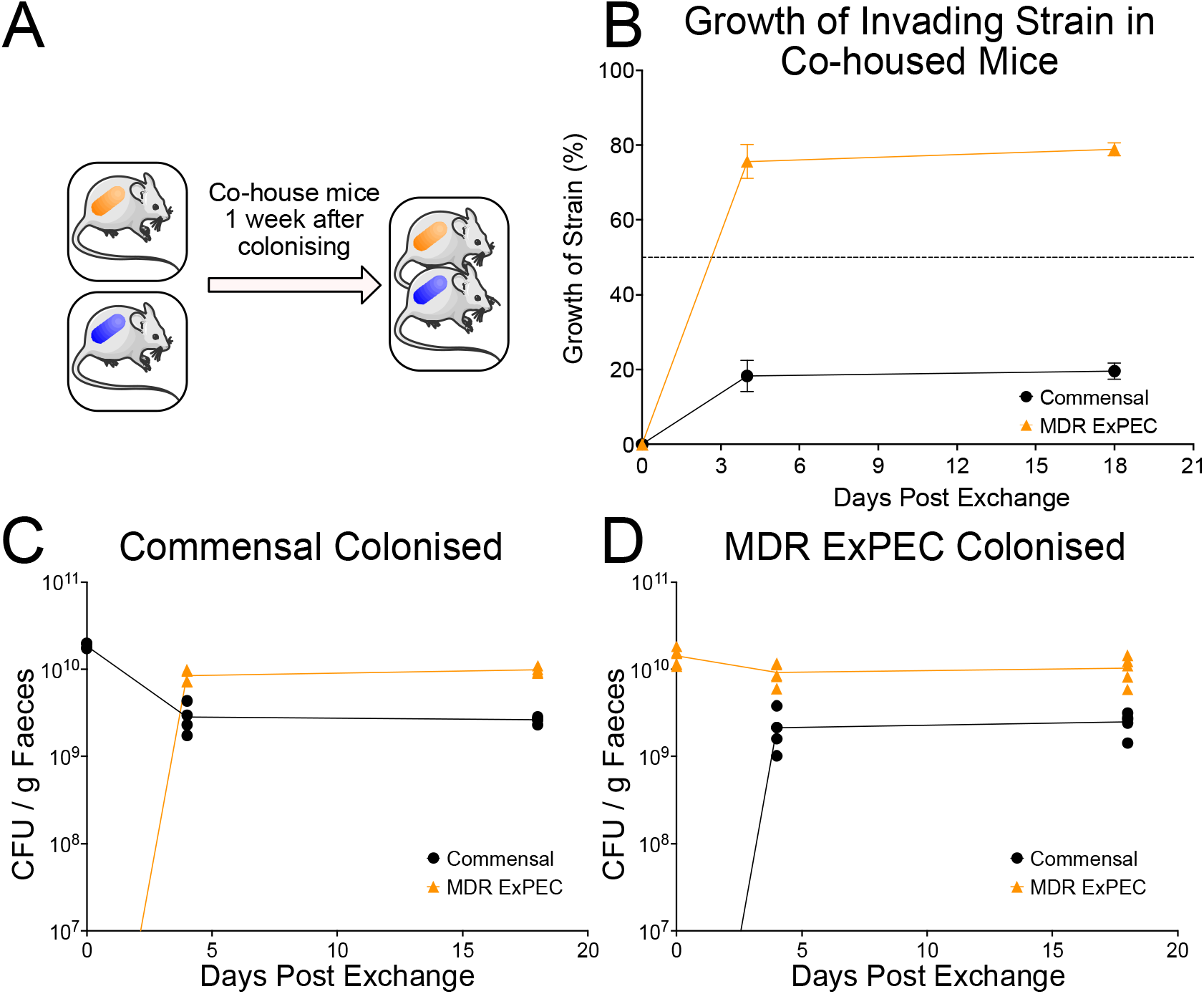
Transmission of strains between monocolonised mice when co-housed. A, Schematic of co-housing experiment in which mice are monocolonised with either commensal or MDR ExPEC strains for one week before exchanging mice between cages. B, Growth of the invading strain in co-housed mice measured by strain specific qPCR at select timepoints (n=4 for commensal shown in black, n=5 for MDR ExPEC shown in orange). Data presented as a percentage of total growth. C and D, CFU per gram of Faeces measured by qPCR in monocolonised mice followed by cohousing. Bacterial growth measured by strain specific qPCR from faecal pellets, commensal growth (black circles) and growth of ST131 (orange triangles). C, Mice monocolonised by a commensal strain (n=4). D, Mice monocolonised by MDR ExPEC (n=5).

### MDR *E. coli* colonisation and displacement results in an inflammatory response in the caecum of mice

Next, we aimed to determine whether MDR *E. coli* colonization/displacement elicited a unique host response in the intestinal mucosa. The host immune response in the small intestine and colon of colonised mice was analysed histologically revealing no evidence of inflammatory cell recruitment or tissue damage in response to MDR ExPEC colonization (Fig 4A, Fig S4 & S5). Expression of 11 cytokines was assayed by probe based qPCR from tissue collected from the small intestine, caecum and colon. Expression of inflammatory cytokines *Tnfa, Cxcl1, Il1b, Il17*, and *S100a8* were all significantly elevated in the caecum of commensally colonised mice 3 weeks after challenge with an MDR ExPEC compared to mice challenged with Phosphate Buffered Saline (PBS) (Fig 4B). This was a delayed host response that was not present 24 hours following challenge (Fig 4C). Interestingly, this host inflammatory response was unique to oral gavage challenge being absent in monocolonized mice 3 weeks after co-housing (Fig 4D). Expression of the assayed cytokines in the small intestine and colon revealed few differences between colonisation conditions (Fig S6, S7 & S8). Collectively, these data demonstrate that displacement of commensal *E. coli* and dominant colonization by MDR ExPEC is associated with a distinct host mucosal immune response.

**Figure 4.**
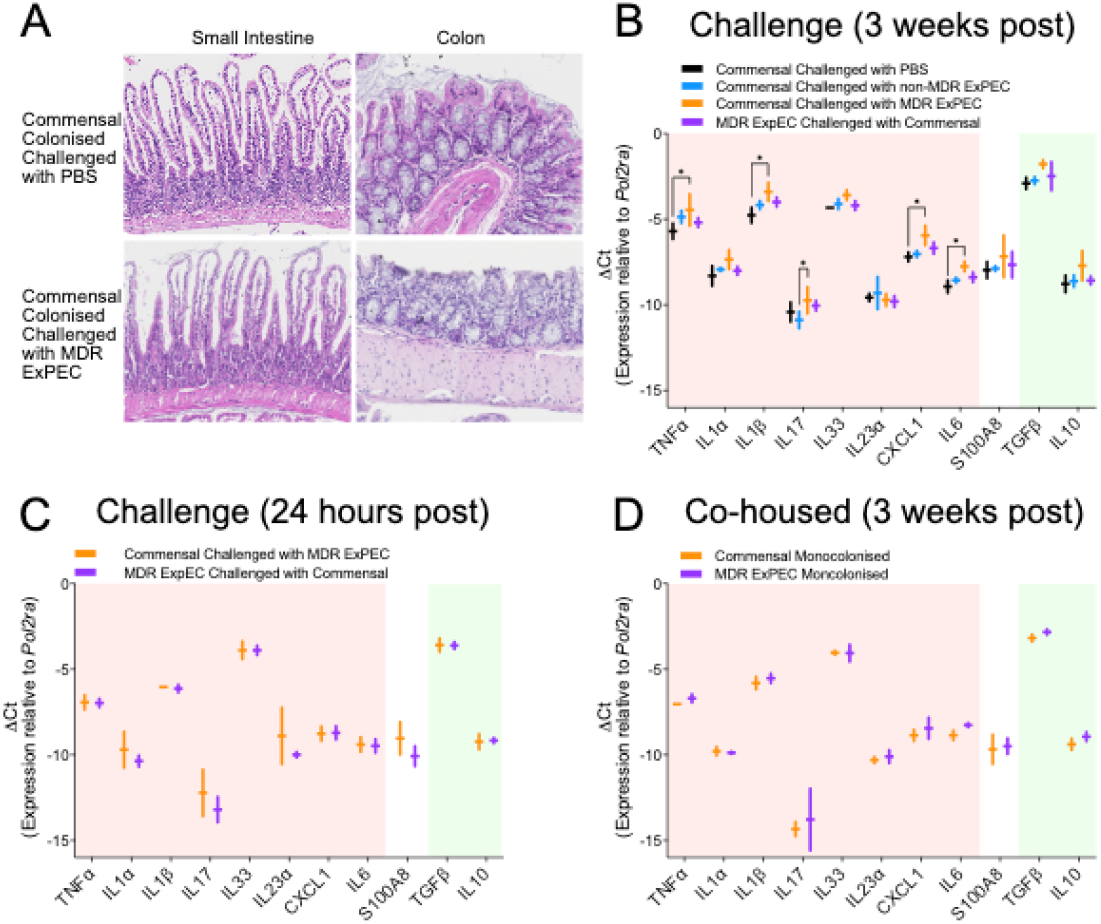
Mouse host inflammatory response to colonisation by multiple strains of *E. coli*. A, Representative histological sections of small intestine and colon from colonised mice, no signs of inflammation were observed under any conditions. B, C, D, Cytokine expression measured by probe based qPCR for a panel of 11 cytokines normalised to endogenous control *Pol2ra*, negative values indicate that level of detected mRNA was less than *Pol2ra* mRNA levels. Cytokines considered to be pro-inflammatory are highlighted in red whilst anti-inflammatory cytokines are in green. B, Expression of assayed cytokines in monocolonised mice 3 weeks after challenge with a second strain. In black commensal monocolonised mice challenged with PBS, in blue commensal monocolonised mice challenged with non-MDR ExPEC, in orange commensal monocolonised mice challenged with MDR ExPEC and in purple MDR ExPEC colonised mice challenged with commensal. C, Expression of assayed cytokines in monocolonised mice 24 hours after challenge with a second strain. In orange commensal monocolonised mice challenged with MDR ExPEC and in purple MDR ExPEC monocolonised mice challenged with commensal. D, Expression of assayed cytokines in monocolonised mice 3 weeks after co-housing. In orange commensal monocolonised mice and in purple MDR ExPEC monocolonised mice. Significance determined by 2-way ANOVA with Tukey’s multiple correction.

Given that our displacement findings cannot be driven by classical virulence genes (Fig S1) we sought to perform a comprehensive genomic analysis of the *E. coli* species to uncover potential genetic factors behind why MDR strains are so dominant in the intestinal tract.

### A curated 20k genome dataset confirms that ESBL and carbapenemase gene carriage is concentrated in a small number of *E. coli* lineages

We curated a dataset of 19,571 *E. coli* genome assemblies encompassing the major *E. coli* phylogroups (A, B1, B2, D, E and F/G), representing 20 STs incorporating commensal, ExPEC and EPEC/EHEC lineages (Figure 5A and Table S3). Antibiotic resistance genes are concentrated in 5 ExPEC lineages: ST38, ST69, ST131, ST167 and ST648 (Figure 5B). These lineages have a high proportion of their population carrying multiple (>1) resistance genes (ST38: 89.0%, ST69: 80.3%, ST131: 85.7%, ST167: 99.1%, ST648: 93.5%) (Fig 5B). Incidence of Beta-lactamase resistance genes is strongly correlated with incidence of resistance genes of any class as a proportion of the lineage (R2=0.935 and P value=4.07e-12). The identified MDR lineages do not cluster on the phylogenetic tree instead spanning multiple phylogroups (Fig 5A).

**Figure 5.**
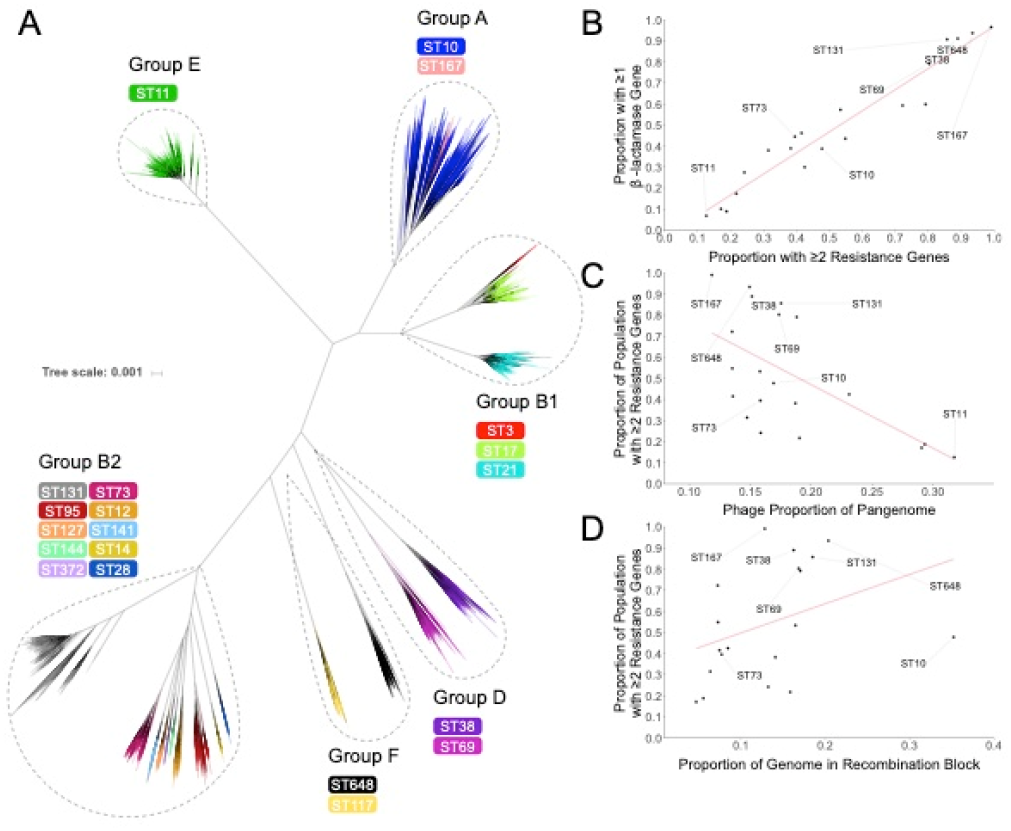
Phylogenetic tree of selected lineages of *E. coli*, incidence of AMR genes within the dataset and correlations between HGT and AMR gene content. A, Phylogenetic tree of dataset produced from Mash distances, tip branches are coloured by ST and phylogroups are outlined. B, The proportion of the lineage with 2 or more resistance genes of any class against the proportion of the lineage with 1 or more beta-lactamase gene. C, Proportion of the pangenome that is associated with phage elements against the proportion of the lineage that has 2 or more resistance genes of any class. D, Proportion of the genome predicted to be within a recombination block against the proportion of the lineage that has 2 or more resistance genes of any class. Red lines indicate linear regressions (B: R^2^ 0.954, P value 1.61 x10^-13^ C: R^2^ 0.369, P value 0.0045, D: R^2^ 0.138, P value 0.117).

Horizontal gene transfer (HGT) is the primary mechanism by which *E. coli* acquires antimicrobial resistance genes such as beta-lactamases. Whilst it is not possible to reliably profile plasmids from the short read data used in this study, we investigated the presence of phage elements in the pangenome alongside recombination. Incredibly AMR carriage is negatively correlated with the proportion of the pangenome associated with phage elements (Fig 5C), whilst AMR carriage is positively correlated with proportion of the genome predicted to be in a recombination block (Fig 5D), meaning that AMR lineages are more recombinogenic but to the exclusion of phage acquisition and mobilisation.

### MDR lineages display an increase in genetic diversity of carbohydrate metabolism genes in their accessory pangenome associated with recombination of new variant alleles

Pangenomes for 20 different lineages of *E. coli* were constructed with Roary v3.10.2 using an identity threshold of 95%, a core gene frequency of 99% with paralog splitting disabled. This setting allows us to specifically look for unique alleles of core genes as those unique alleles then become part of the accessory genome^13^. The host generalist ST10 had the greatest pangenome size with 46,259 gene clusters identified followed by ST131 with 23,857 clusters (Table S4, Figure S14). Core genome size was consistent across all lineages averaging 3777 genes (Table S4, Figure S15). There was no correlation between pangenome size and carriage of AMR genes (Fig S13). Pangenomes were functionally annotated using EggNOG database with the eMapper utility and functional composition of the pangenome was explored using Clusters of Orthologous Groups (COG) categories. Links between AMR and biological function were explored in the core and accessory genome using linear regressions analysis which revealed a single significant association. Specifically, lineages with a higher proportion of AMR genes displayed an increased number of ‘Carbohydrate metabolism and transport’ genes in their accessory genome (Fig 6A). There were no other significant correlations between COG categories and carriage of AMR genes after correcting for multiple testing. Our data suggest that diversification of carbohydrate metabolic genes is correlated to the acquisition of multi-drug resistance in *E. coli*.

**Figure 6.**
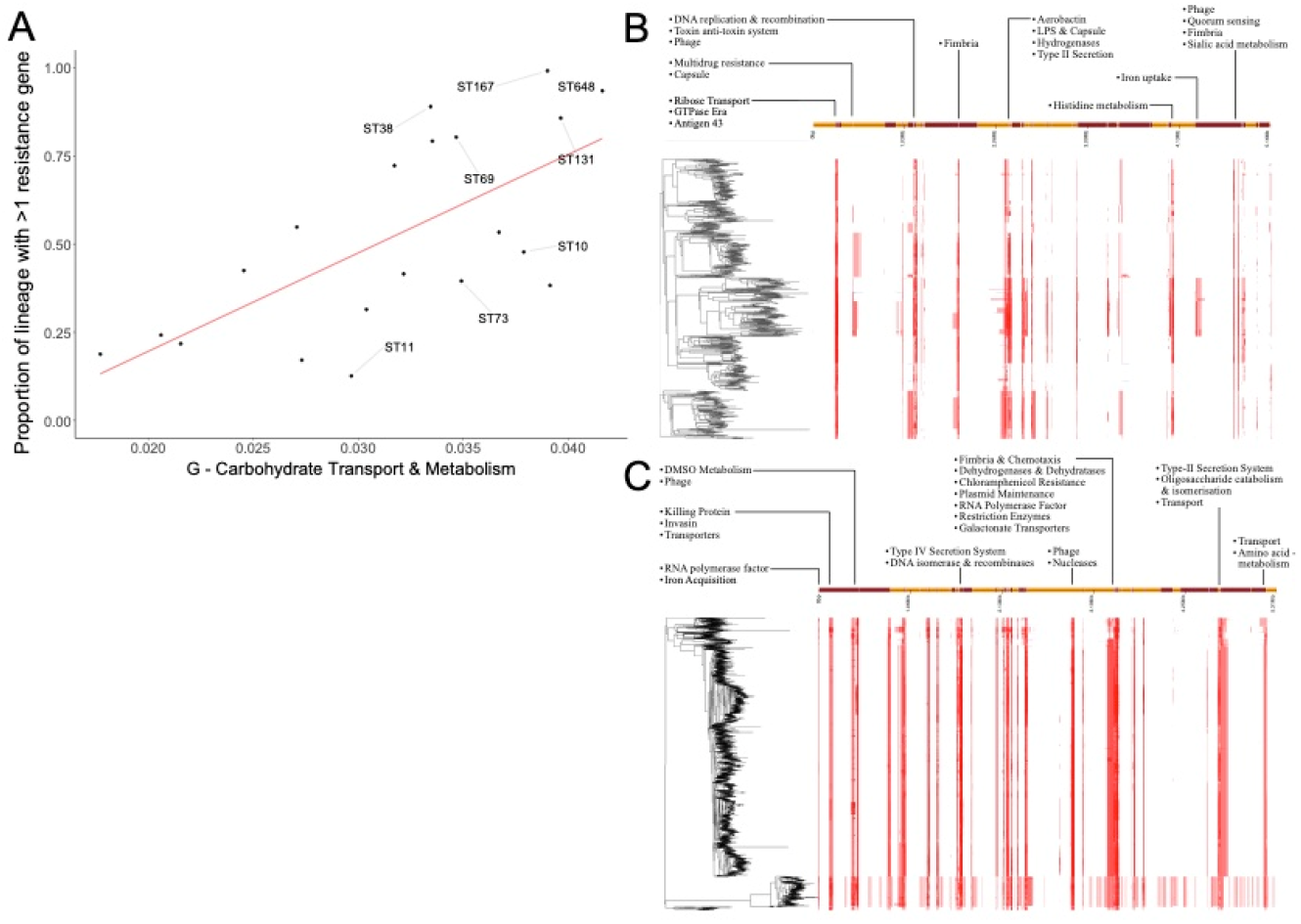
A. The proportion of the accessory pangenome annotated with G COG category (carbohydrate transport and metabolism) against the proportion of each lineage with more than 2 resistance genes. Red line indicates linear regression line. (R-squared of 0.4518, P value of 0.0007). B,C. Recombination regions predicted by Gubbins in ST73 (B) and ST131 (C) lineages.

We sought to determine the source driving our observed diversity of carbohydrate genes AMR lineages. Our analysis confirmed that the metabolism genes being identified in the accessory genome were in fact allelic variants of genes core to *E. coli*. We then looked at the recombination plot created for the main MDR *E. coli* lineage ST131 (Fig 6B) and the main non-MDR ExPEC ST73 (Fig 6C). Our analysis clearly shows recombination occurring in key metabolic loci that does not occur in ST73 and that these loci house metabolism genes occurring as allelic variants in the accessory genome dataset.

## Discussion

The evolution and global transmission of AMR are a major threat to public health. Genomic identification of antibiotic resistance genes in our global dataset highlights that AMR is concentrated in a number pandemic lineages. This is most evident when examining the average number of resistance genes per genome; ST167, ST648, ST38 and ST131 all display on average in excess of 7 resistance genes per genome. This result is not driven by many resistance genes within a small subpopulation, as ST167, ST648, ST38 and ST131 all have in excess of 80% of their population possessing multiple resistance genes highlighting the success of these lineages. This observation is aligned with numerous other studies which frequently report these populations as major MDR pathogens, confirming previous observations that AMR carriage is not equal across the *E. coli* population with AMR being concentrated in certain lineages^2^.

Successful pandemic clones must transmit from the environment to individuals, or between individuals rapidly in order to spread globally. Here we demonstrate that an MDR ST131 strain can readily colonise new hosts even when those hosts are pre-colonised with commensal *E. coli*. The invading ST131 becomes the dominant colonising strain in mice, both when the invading strain is introduced artificially via oral gavage and when mice are co-housed. It is important to emphasise that this transmission is occurring in the absence of any antibiotic treatment conferring a selective advantage to ST131. Typically it has been required that mice are treated with streptomycin to allow *E. coli* strains to colonise them, however more recent studies, alongside data presented here, indicate that ST131 strains do not require antibiotic treatment in order to competitively colonise mice^27^. In our study the ST131 out-colonises another ExPEC strain of the common ST73 lineage. This implies that ST131 possesses some mechanism by which it can out-compete commensals that is lacking in another highly successful but non-MDR ExPEC lineage ST73. Our observations from mouse colonisation are supported by previous studies which have observed MDR *E. coli* as frequent colonisers of healthy travellers in the absence of antibiotics^7–9^. Within household transmission has also been observed for ST131^28^. Studies of healthy individuals who travel to regions where antibiotic resistance is endemic have reported varying levels of colonisation of between 30% and 70% upon return^7^. Sampling travellers during their trip revealed a much higher rate of colonisation rate of 95%, of which *E. coli* was the most common colonising bacteria^9^. From our data the resident commensal strain was not completely displaced, similar observations have been made of human travellers, specifically individuals colonised by MDR *E. coli* had a recurrence of their original commensal *E. coli* at the end of their travels^8^.

Three weeks after mice were challenged with ST131 we detected increased mRNA levels of the pro-inflammatory cytokines TNFα, IL1β, IL-6 and CXCL1 in the caecum. Expression of TNFα by plasma cells has been implicated in pathogen clearance in mice^29^, while IL-1β is only produced by mononuclear phagocytes upon exposure to pathogens and not commensals^30^. IL-1β has also been implicated as important for repair of the epithelial barrier following colitis^31^. CXCL1 is released from epithelial cells to recruit neutrophils to aid in pathogen clearance^32^, lastly IL-6 has diverse pro-inflammatory effects including effector cell differentiation^33^. Together the upregulation of these cytokines suggest that the host is responding to MDR *E. coli* ST131 as a pathogen and that it is attempting to clear it from the intestine. Our observed increase in inflammatory cytokines is confined to the caecum, whilst the other assayed tissues displayed alterations in cytokine expression that were not unique to ST131. Previous work has demonstrated that ST131 colonisation burden is highest in the caecum compared to other compartments^27^, an observation we also make. There have been limited investigations into the human immune response to colonisation by MDR bacteria however it has been reported that traveller’s diarrhoea is the most frequently reported symptom by those who are colonised^34^, suggesting that there may be a mild inflammatory response in humans following colonisation. What is striking is that our observations are similar to those made for the initial colonisation of pathogenic *Salmonella* and *E. coli* in the mammalian gut, whereby localised inflammation facilitates outcompetition of commensal bacteria by the invading pathogen^35–37^.

To identify the underlying genetic mechanisms underlying MDR pandemic *E. coli* clones ability to dominantly colonize the mammalian gut, we used a functional pangenome analysis pipeline. Previous pangenome analysis identified metabolic loci as being enriched in nucleotide diversity in ST131 compared to other ExPEC, specifically anaerobic metabolic genes were exhibiting increased genetic variation^13^. Our functional pangenome analysis supports these observations, revealing that there is a significant correlation between carriage of AMR genes and genetic diversity in metabolism genes. Specifically, there is increased variation in genes encoding carbohydrate metabolism in lineages with a high rate of antibiotic resistance carriage. Previous analyses have focussed on individual lineages whereas here we present data on multiple MDR ExPEC lineages, revealing that metabolic variation is a shared adaptation of MDR *E. coli*. Together these point towards a pivotal role for metabolism in MDR lineage formation. Experimental evolution studies have identified that *E. coli* can evolve resistance to antibiotic stress through mutations in core metabolic genes, particularly those involved in carbon and energy metabolism^38^. These observed mutations occur at low frequency and were only detected through sequencing of multiple isolates from a population; however, they were still detectable in datasets of clinical samples demonstrating their relevance. A large scale bacterial genetic screen to test the effect of allelic variation in key metabolism genes on the ability to colonise and displace commensal *E. coli* in the mammalian intestinal tract would seem attractive. However the signal observed in our dataset occurs in multiple genes and pathways and genetically investigating such a polygenic trait is far from trivial. We suggest further investigation of our findings will need to combine classical genetics with long term and complex experimental studies attempting to recapitulate MDR clone evolution.

Collectively, our data demonstrate that MDR *E. coli* is highly capable of host intestinal colonisation, displacing resident commensal *E*. coli to become the dominant strain and readily transmitting between hosts. Our pangenomic analysis implicates metabolism as a pivotal factor in the evolution of AMR linked to the incredible gut colonisation ability of MDR *E. coli*.

## Acknowledgements

This works was funded by a Wellcome Trust funded MIDAS PhD studentship awarded to CC (Grant number 203821/Z/16/A).

