## Supplementary files for "Multi-drug resistant *E. coli* displace commensal *E. coli* from the intestinal tract, a trait associated with elevated levels of genetic diversity in carbohydrate metabolism genes"

Table S1: *E. coli* strains used for mouse colonisation experiments

| Strain | ID | ST | Source | Accession | Reference: |
| --- | --- | --- | --- | --- | --- |
| Commensal | 822-E8 | ST73 | Healthy Human Volunteer | PRJNA893850 | Strain a gift from Prof Peter Hawkey |
| Non-MDR ExPEC | F084 | ST73 | Blood Stream Infection | PRJEB9931 | 10.1093/jac/dkv365 |
| MDR ExPEC | F016 | ST131 | Blood Stream Infection | SRR3143583 | 10.1371/journal.pgen.1006280 |

Table S2: primers and probes used for qPCR – see excel spreadsheet

Table S3: Genomes analysed

|  | Sequence Type | Phylogroup | Genomes Downloaded | Genomes Analysed | Comments |
| --- | --- | --- | --- | --- | --- |
| 1 | ST3 | B1 | 40 | 40 | EPEC |
| 2 | ST10 | A | 4090 | 2370 | Generalist |
| 3 | ST11 | E | 6230 | 5137 | EHEC O157 |
| 4 | ST12 | B2 | 311 | 283 | ExPEC |
| 5 | ST14 | B2 | 63 | 62 | ExPEC |
| 6 | ST17 | B1 | 1942 | 1884 | EHEC |
| 7 | ST21 | B1 | 2504 | 2411 | EHEC |
| 8 | ST28 | B2 | 47 | 46 | EPEC |
| 9 | ST38 | D | 662 | 617 | ExPEC |
| 10 | ST69 | D | 759 | 696 | ExPEC |
| 11 | ST73 | B2 | 946 | 873 | ExPEC |
| 12 | ST95 | B2 | 805 | 758 | ExPEC |
| 13 | ST117 | F | 322 | 269 | ExPEC |
| 14 | ST127 | B2 | 241 | 232 | ExPEC |
| 15 | ST131 | B2 | 3772 | 3186 | ExPEC |
| 16 | ST141 | B2 | 94 | 91 | ExPEC |
| 17 | ST144 | B2 | 65 | 65 | ExPEC |
| 18 | ST167 | A | 117 | 115 | ExPEC |
| 19 | ST372 | B2 | 56 | 54 | ExPEC |
| 20 | ST648 | F | 450 | 382 | ExPEC |
|  | Total |  | 23516 | 19571 |  |

Table S4: Pangenome sizes

|  | Sequence Type | Phylogroup | Genomes Analysed | Comments | Pangenome Size | Core Genome Size |
| --- | --- | --- | --- | --- | --- | --- |
| 1 | ST3 | B1 | 40 | EPEC | 8417 | 3716 |
| 2 | ST10 | A | 2370 | Generalist | 46259 | 3104 |
| 3 | ST11 | E | 5137 | EHEC O157 | 20997 | 3990 |
| 4 | ST12 | B2 | 283 | ExPEC | 13876 | 3786 |
| 5 | ST14 | B2 | 62 | ExPEC | 13876 | 3853 |
| 6 | ST17 | B1 | 1884 | EHEC | 16370 | 4049 |
| 7 | ST21 | B1 | 2411 | EHEC | 14634 | 4160 |
| 8 | ST28 | B2 | 46 | EPEC | 8796 | 3603 |
| 9 | ST38 | D | 617 | ExPEC | 20765 | 3757 |
| 10 | ST69 | D | 696 | ExPEC | 21345 | 3693 |
| 11 | ST73 | B2 | 873 | ExPEC | 16431 | 3868 |
| 12 | ST95 | B2 | 758 | ExPEC | 16677 | 3825 |
| 13 | ST117 | F | 269 | ExPEC | 14582 | 3792 |
| 14 | ST127 | B2 | 232 | ExPEC | 12260 | 3850 |
| 15 | ST131 | B2 | 3186 | ExPEC | 23857 | 3641 |
| 16 | ST141 | B2 | 91 | ExPEC | 10479 | 3856 |
| 17 | ST144 | B2 | 65 | ExPEC | 8307 | 3867 |
| 18 | ST167 | A | 115 | ExPEC | 11704 | 3666 |
| 19 | ST372 | B2 | 54 | ExPEC | 8639 | 3737 |
| 20 | ST648 | F | 382 | ExPEC | 15130 | 3736 |

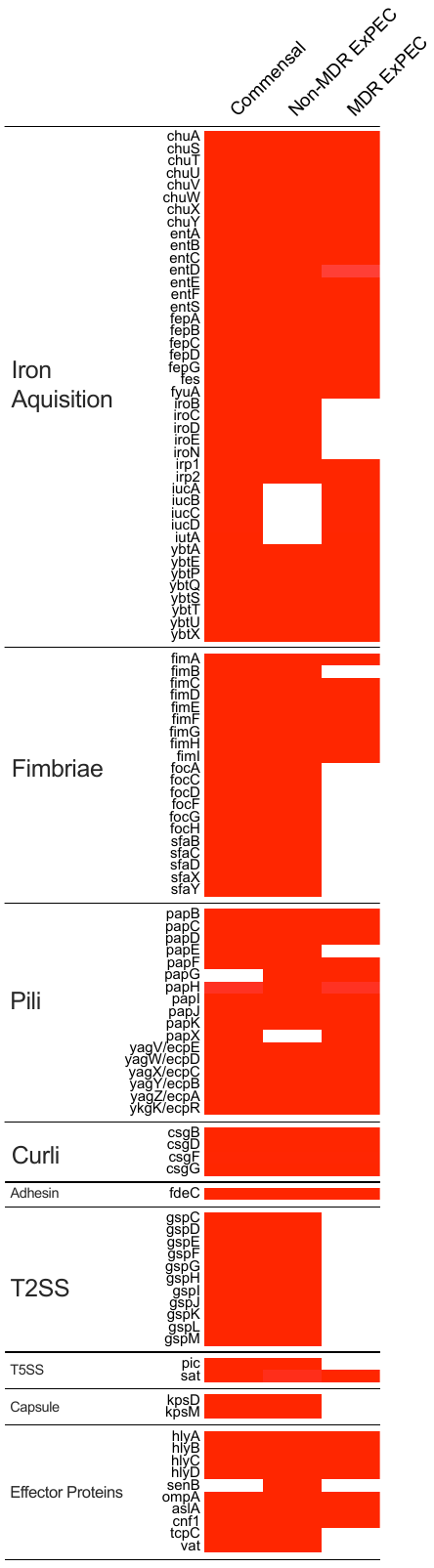

Figure S1: Heatmap of virulence factors identified in mouse colonising strains

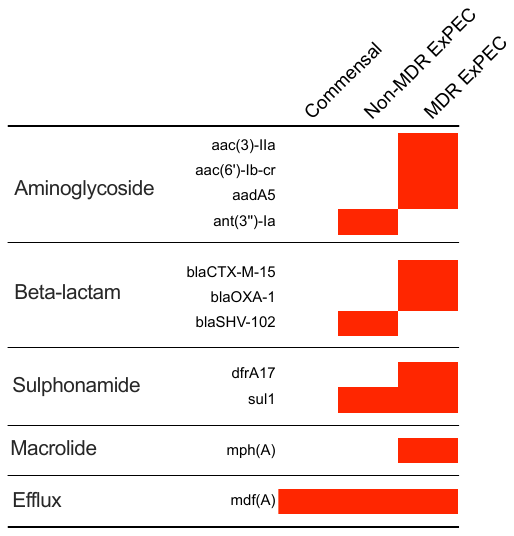

Figure S2: Heatmap of AMR genes identified in mouse colonising strains

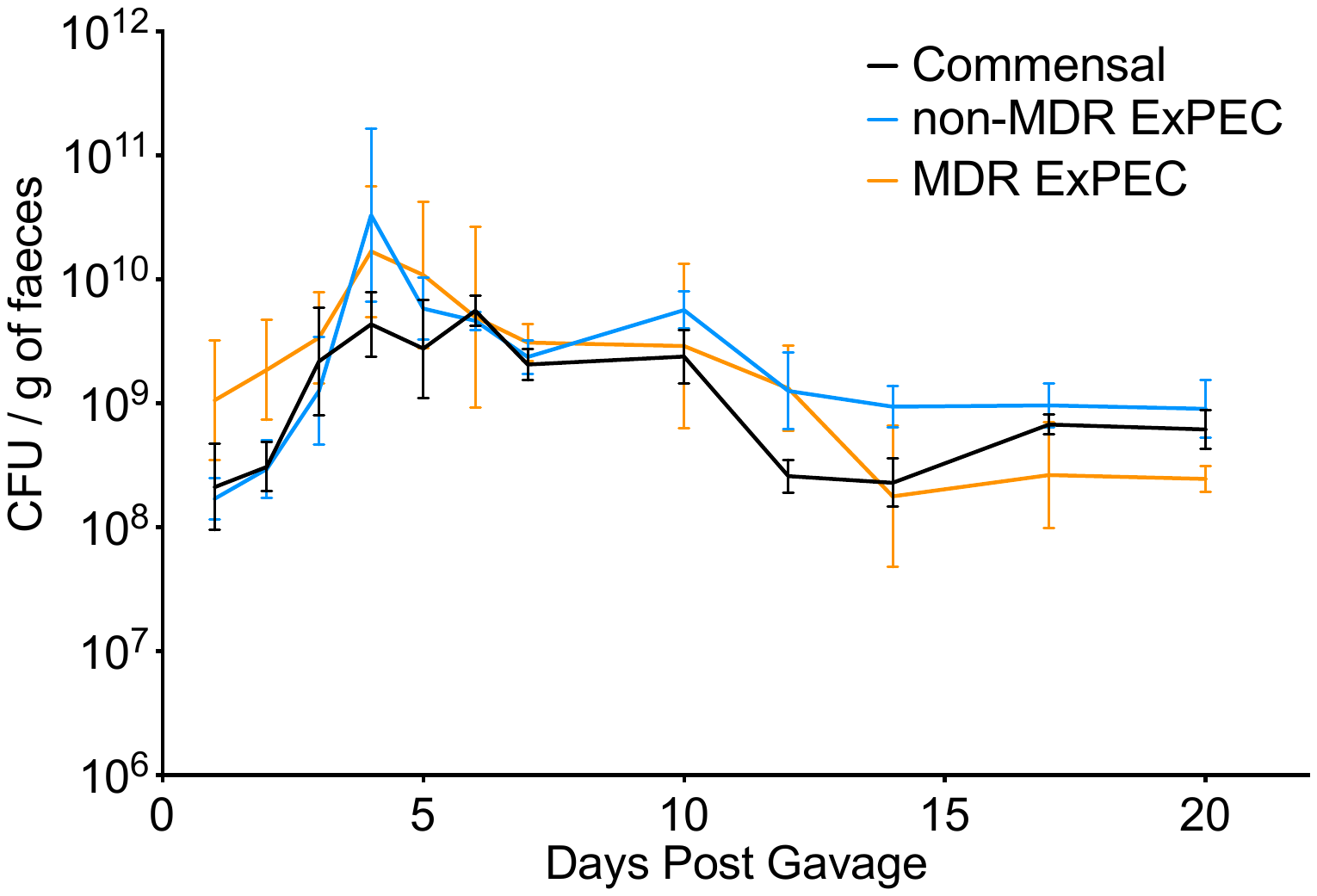

Figure S3: CFU of colonising bacteria in monocolonised mice measured by enumeration from UTI Chromogenic agar plates. CFU counts were adjusted to faecael pellet weight. Geometric average CFU of Commensal (black line), non-MDR ExPEC (blue line) and MDR ExPEC (organge line) is shown with geometric standard deviation, 3 replicates for each condition.

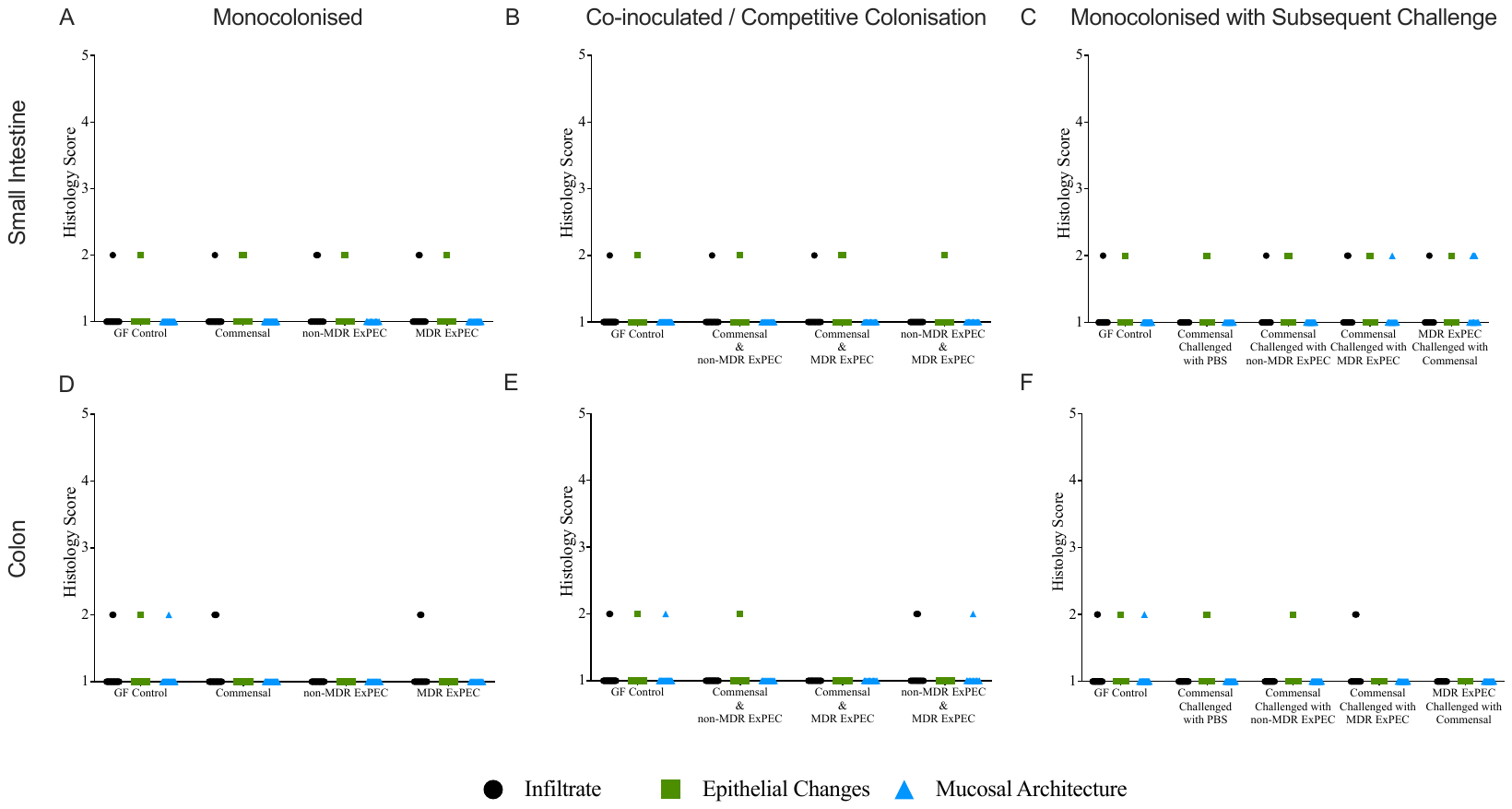

Figure S4: Inflammatory histology score for sections of small intestine and colon from colonised mice and germ free control mice. Small intestine histology scores for monocolonised mice (A), co-incolucated / competitive colonisation mice (B) and monocolonised mice challenged with a second strain after 1 week (C). Colon section histology scores for monocolonised mice (D), co-incolucated / competitive colonisation mice (E) and monocolonised mice challenged with a second strain after 1 week (F). Scores were assigned to inflammatory cell infiltrate (black circles), changes to epithelium (green squares) and overall mucosal architecture (blue triangles). Score of 1 indicates homeostatic mucosa while a score of 5 indicates severe inflammatory disruption. Points represent scores for individual fields of view (FoV) taken from multiple tissue sections.

Number of fields of view scored for each condition

| Colonisation Condition | | Small Intestine | Colon |
| --- | --- | --- | --- |
| Monocolonised | Commensal (n=3) | 26 | 19 |
|  | non-MDR ExPEC (n=3) | 12 | 18 |
|  | MDR ExPEC (n=3) | 19 | 15 |
| Co-inoculated / Competitive Colonisation | Commensal & non-MDR ExPEC | 15 | 12 |
|  | Commensal & MDR ExPEC | 12 | 12 |
|  | Non-MDR ExPEC & MDR ExPEC | 12 | 12 |
| Monocolonised with challenge after 7 days | Commensal challenged with PBS | 12 | 15 |
|  | Commensal challenged with non-MDR ExPEC | 15 | 15 |
|  | Commensal challenged with MDR ExPEC | 12 | 12 |
|  | MDR ExPEC challenged with commensal | 12 | 12 |
| Germ Free | Germ Free | 21 | 21 |

Figure S5: Representative histological sections of colonised mouse small intestine and colon stained with H&E. Germ free control mice are included.

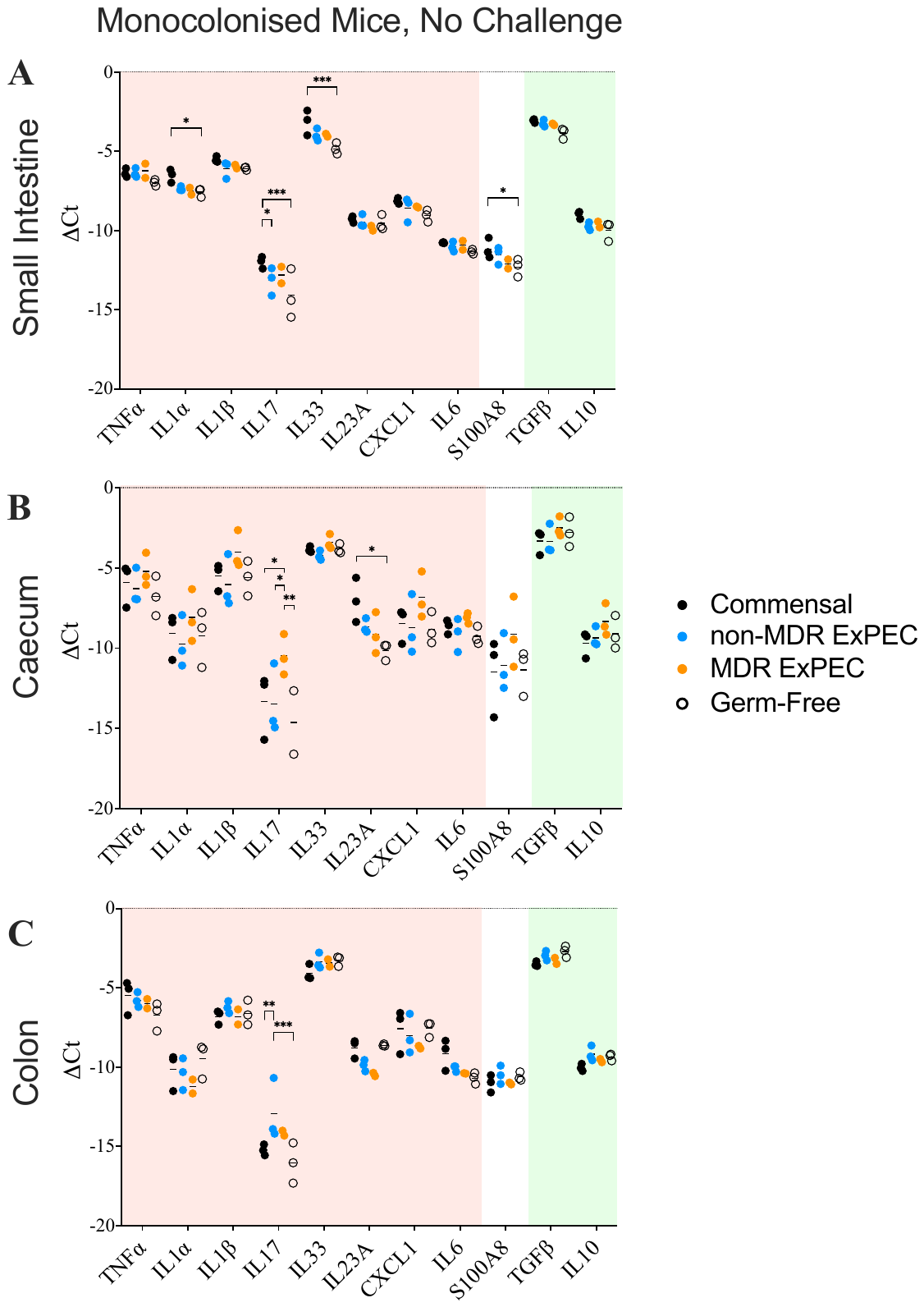

Figure S6: Cytokine expression in the small intestine (A), caecum (B) and colon (C) of monocolonised mice 3 weeks after colonisation. Mice were colonised with a commensal strain (black), non-MDR ExPEC (blue) or MDR ExPEC (orange). Germ free mice were also assayed (open circles). Expression of cytokines is displayed as the delta Ct between cytokine and housekeeping gene Pol2ra, negative values indicate lower expression than housekeeping gene. Statistical significance determined with 2way ANOVA with Tukey’s multiple comparisons, n=3 for all conditions.

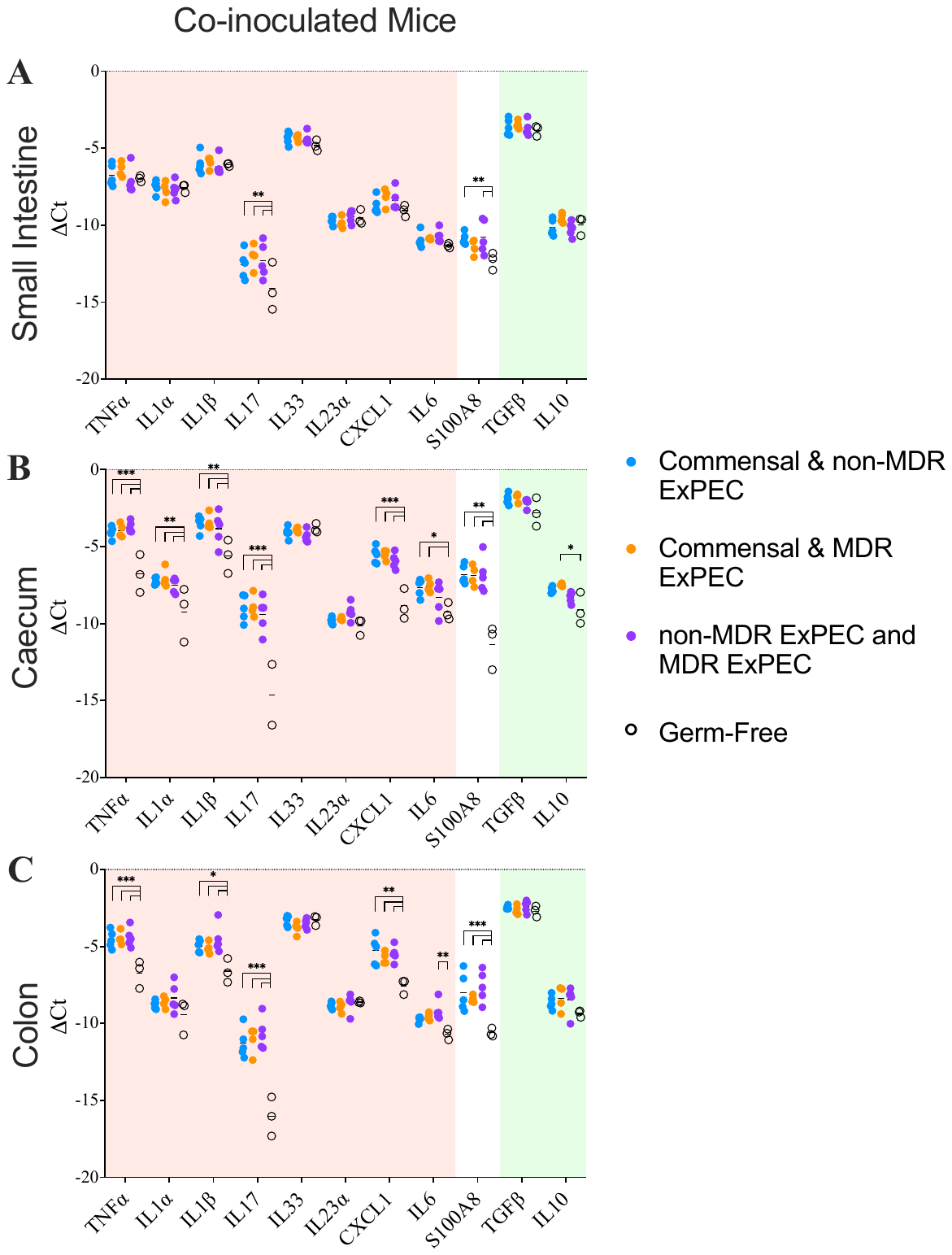

Figure S7: Cytokine expression in the small intestine (A), caecum (B) and colon (C) of co-inoculated mice 1 week after colonisation. Mice were colonised with a commensal strain and non-MDR ExPEC (blue, n=5), a commensal and MDR ExPEC (orange, n=4), or a non-MDR ExPEC and MDR ExPEC (purple, n=5). Germ free mice were also assayed (open circles, n=3). Expression of cytokines is displayed as the delta Ct between cytokine and housekeeping gene Pol2ra, negative values indicate lower expression than housekeeping gene. Statistical significance determined with 2way ANOVA with Tukey’s multiple comparisons.

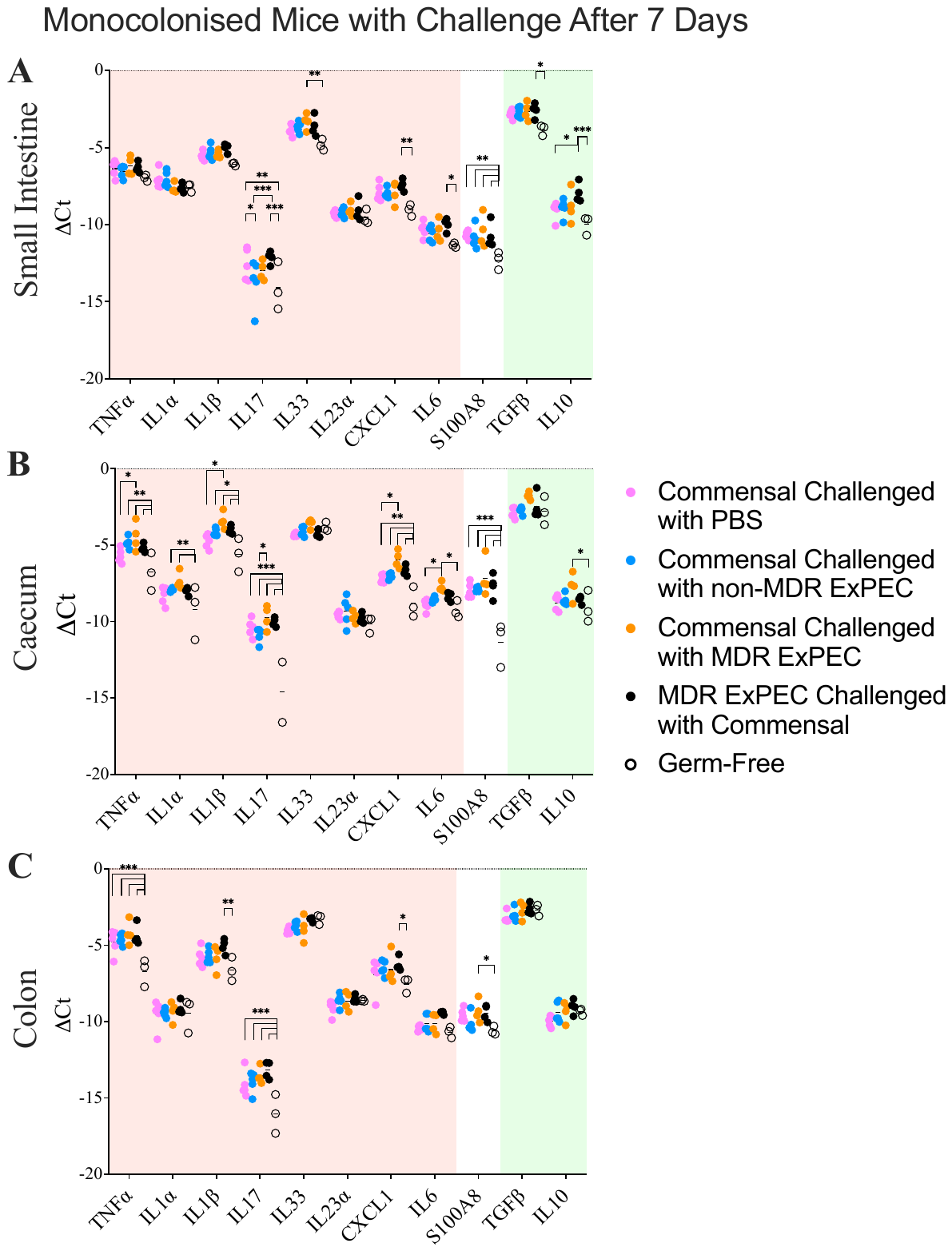

Figure S8: Cytokine expression in the small intestine (A), caecum (B) and colon (C) of monocolonised mice which were challenged with a second strain 7 days after initial colonisation. Mice were colonised with a commensal strain and challenged with PBS (pink, n=5), a commensal strain challenged with a non-MDR ExPEC (blue, n=5), a commensal challenged with an MDR ExPEC (orange, n=4), or an MDR ExPEC challenged with a commensal (black, n=4). Germ free mice were also assayed (open circles, n=3). Expression of cytokines is displayed as the delta Ct between cytokine and housekeeping gene Pol2ra, negative values indicate lower expression than housekeeping gene. Statistical significance determined with 2way ANOVA with Tukey’s multiple comparisons.

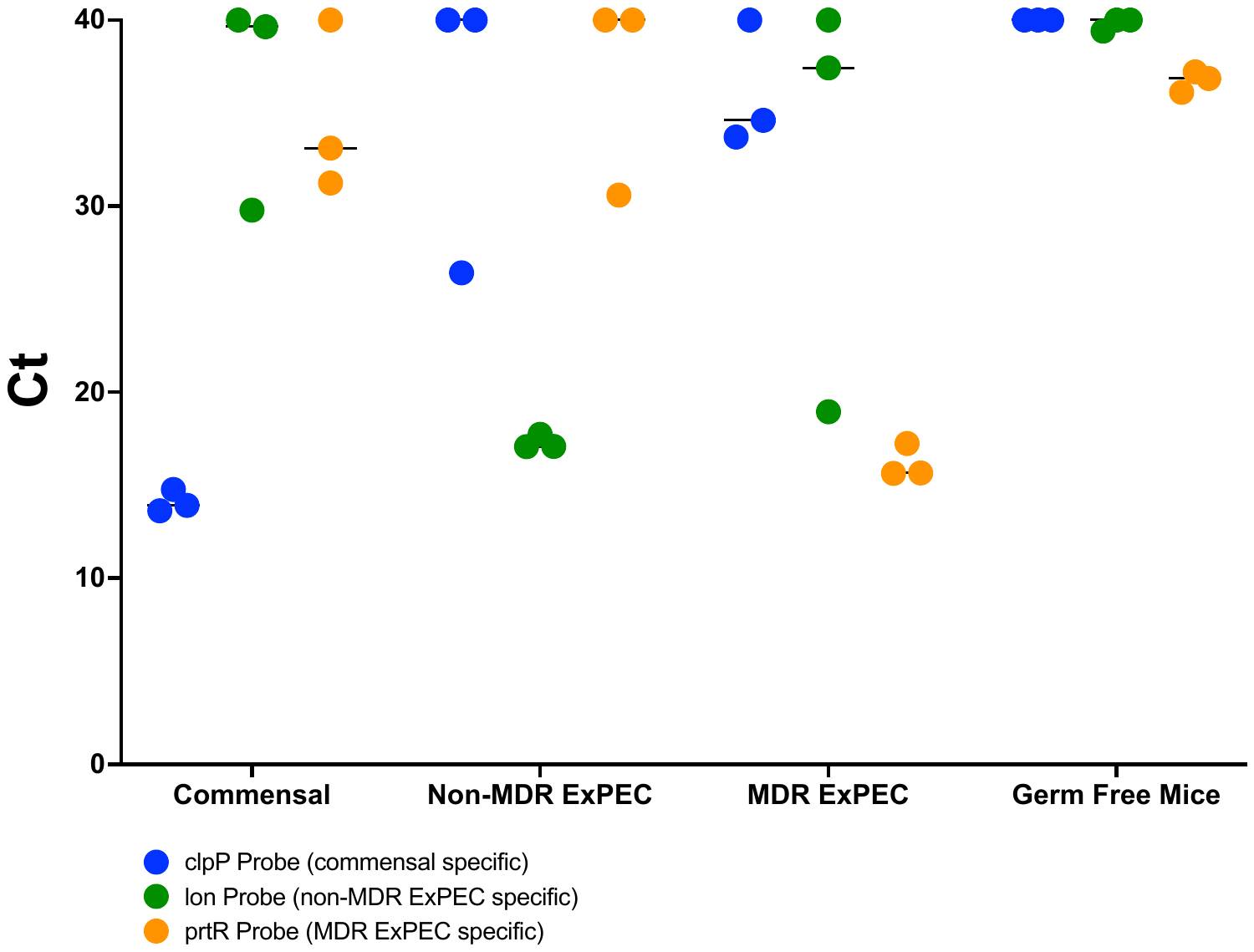

Figure S9: Cycle threshold values for strain specific probes assayed using DNA extracted from faecal pellets from monocolonised mice. Commensal specific probe (clpP) in blue, non-MDR ExPEC probe (lon) in green and MDR ExPEC probe (prtR) in organge.

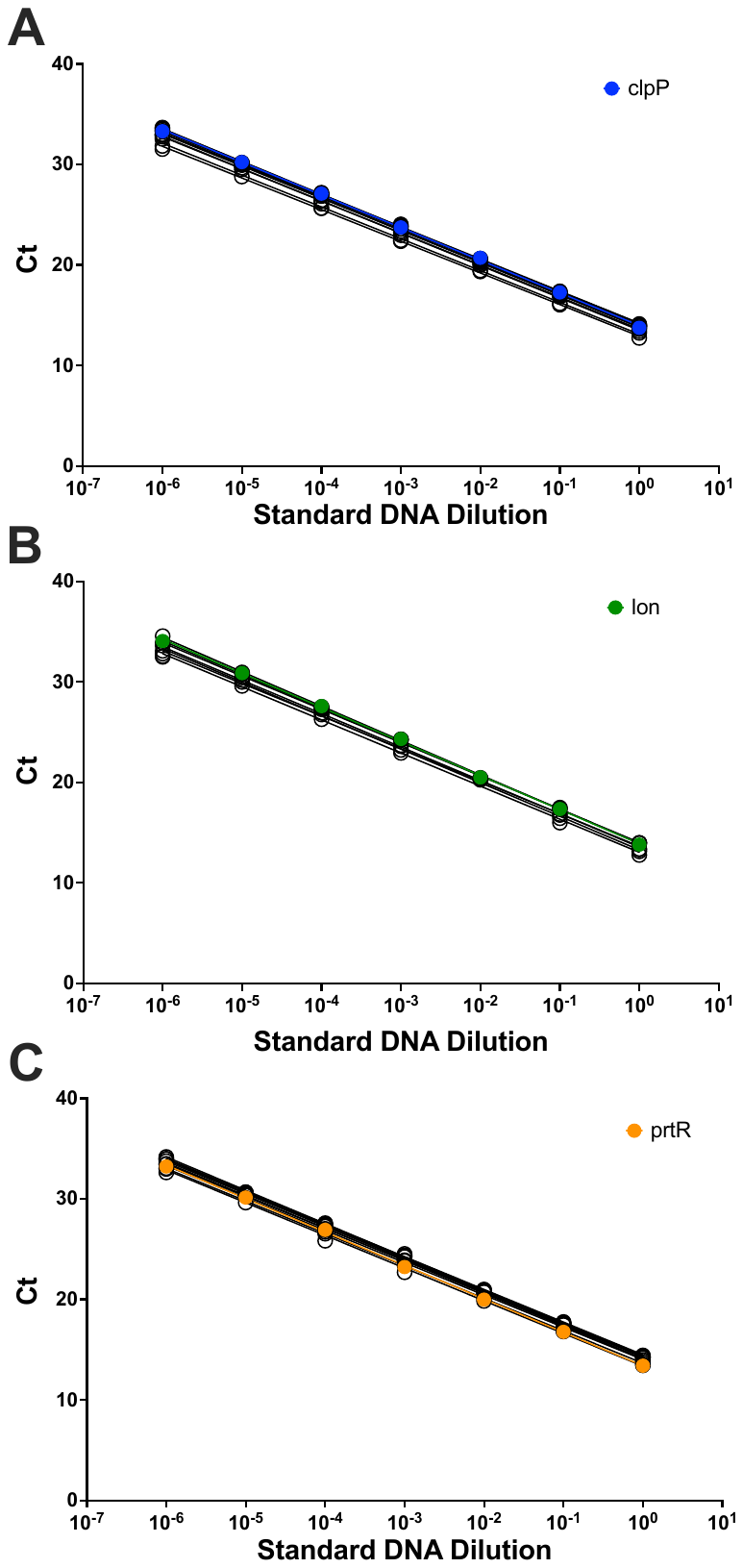

Figure S10: Standard curves for CFU determination. DNA isolates from broth culture of known CFU serially 10 fold diluted and plotted against Ct for that dilution. Colour lines indicate standard curve from stock dilutions while black lines indicate standard curves from every assay plate used for CFU quantitation. Standard curves for commensal specific probe clpP (A), non-MDR ExPEC probe lon (B) and MDR ExPEC probe prtR (C).

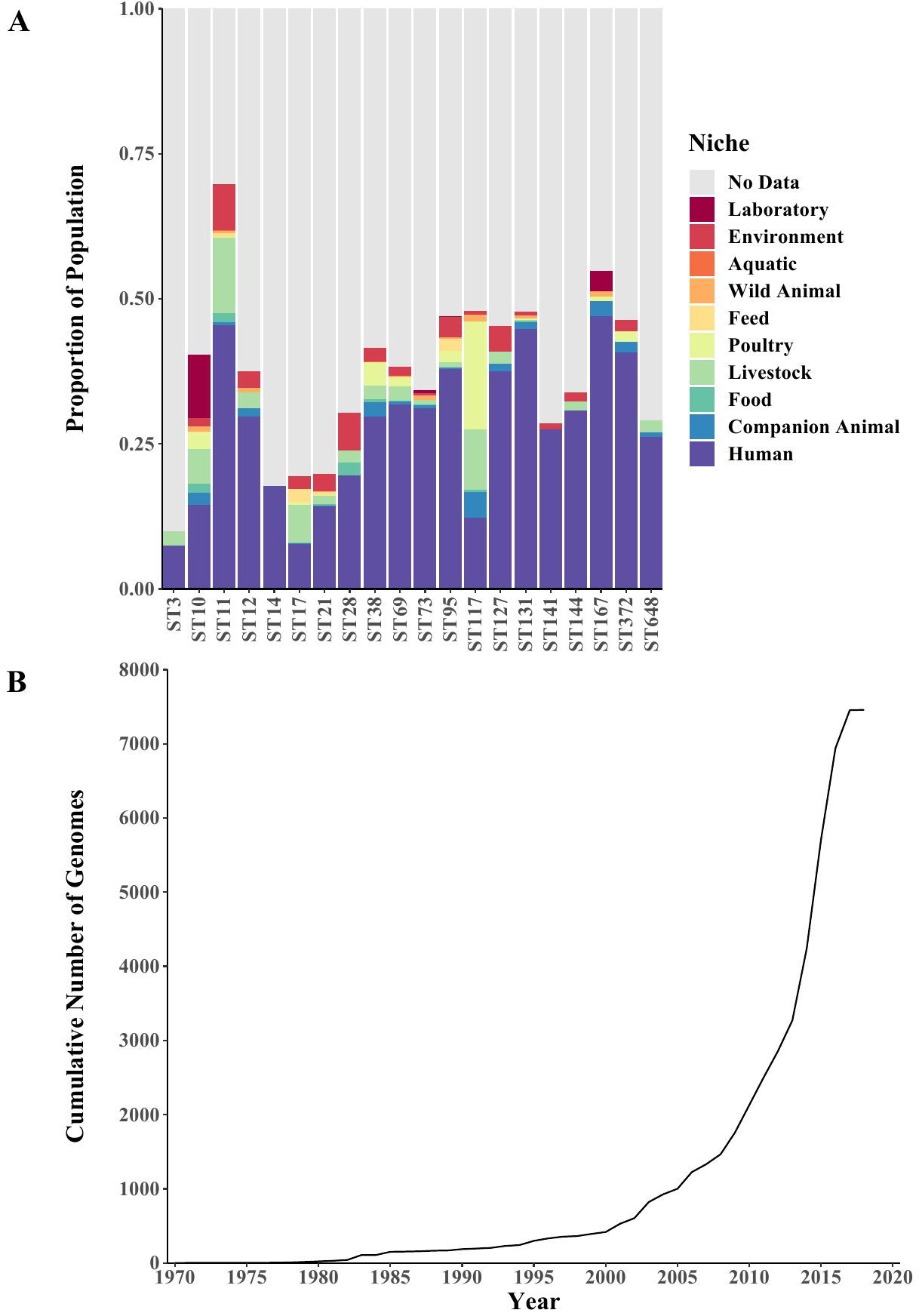

Figure S11: Source niche for genome in dataset (A) and cumulative time of collection (B).

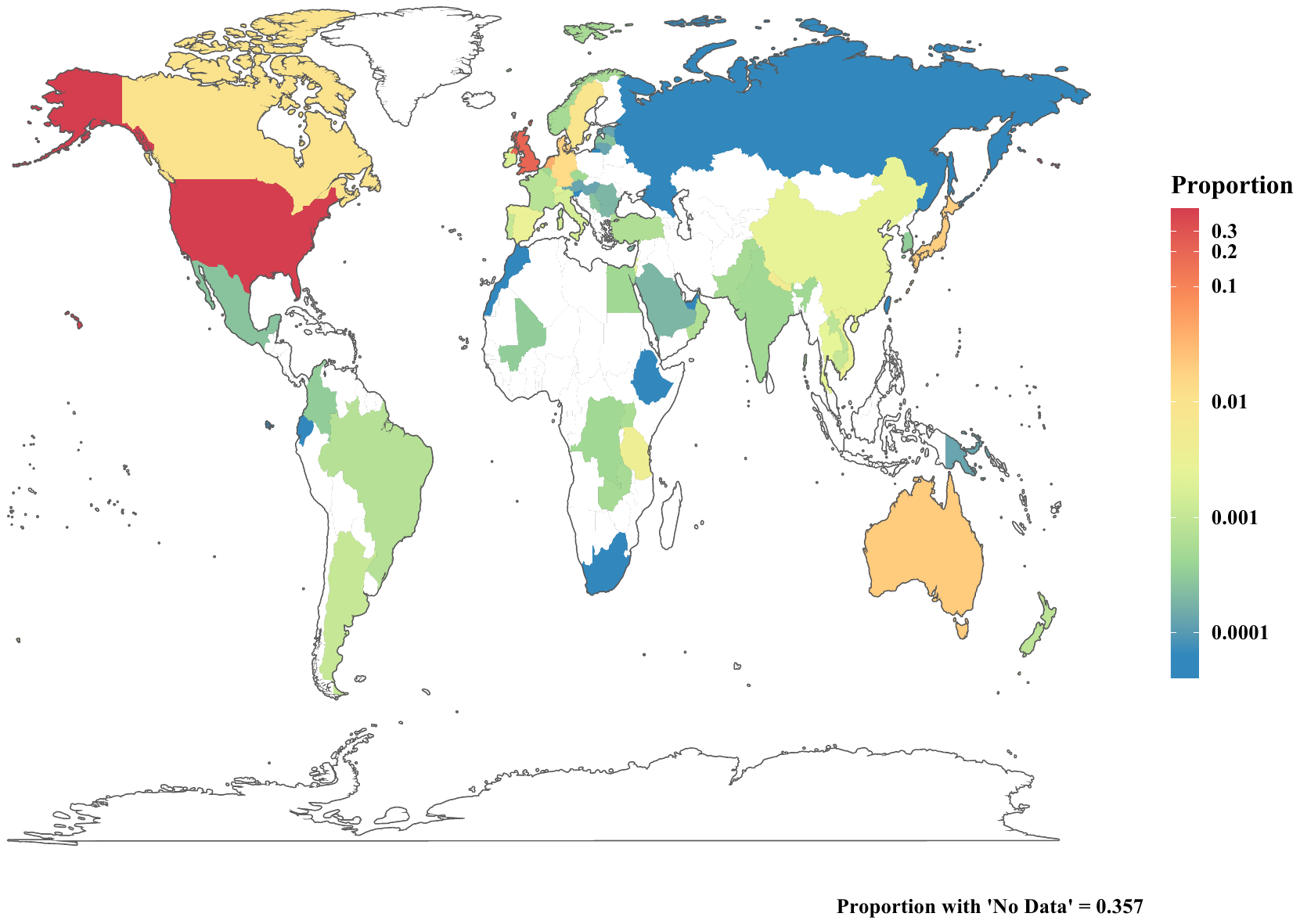

Figure S12. Geographic distribution of genomes in dataset, colour scale is logarithmic.

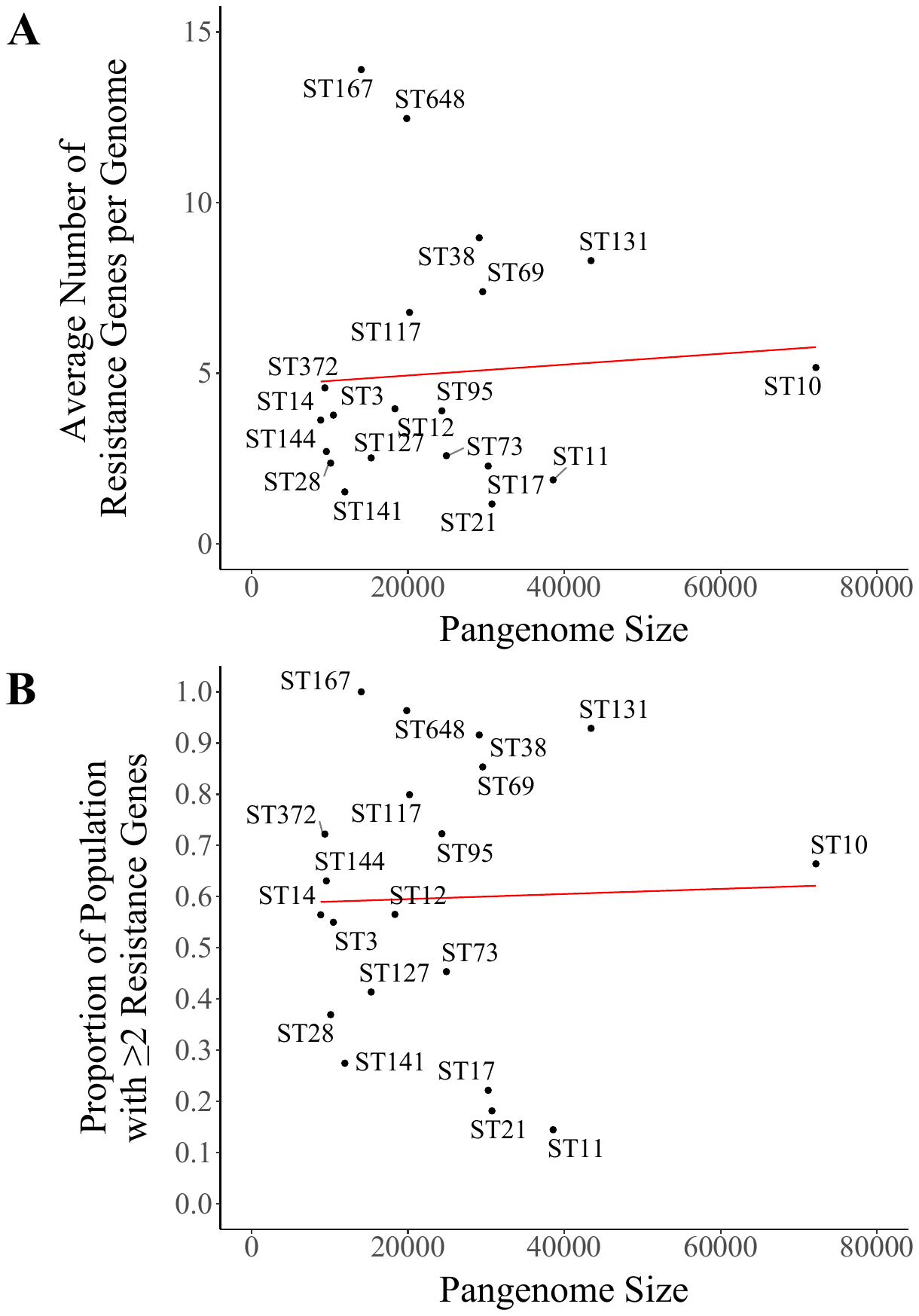

Figure S13: A- Correlation between pangenome size and average number of resistance genes. B- Correlation between pangenome size and proportion of the lineage with 2 or more resistance genes.

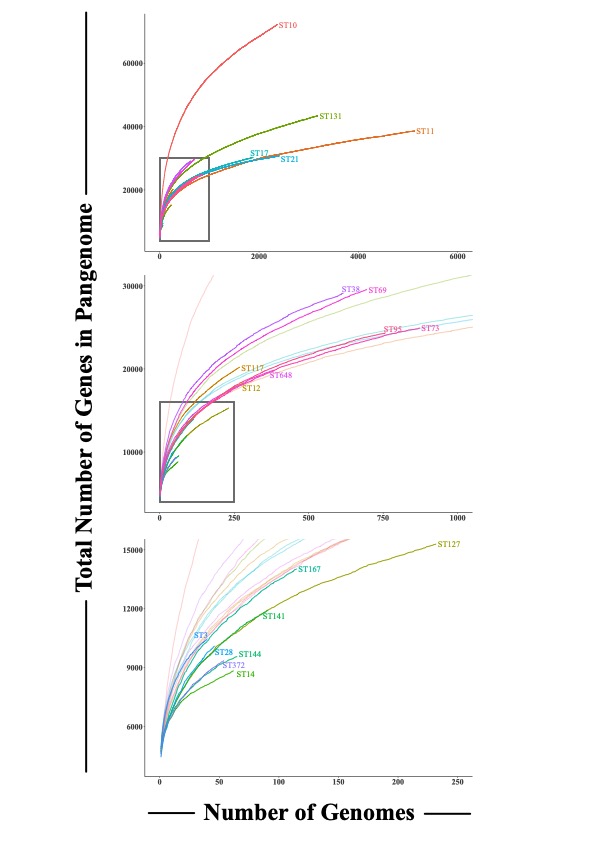

Figure S14: Pangenome rarefaction curves for the accessory compartment of each ST pangenome.

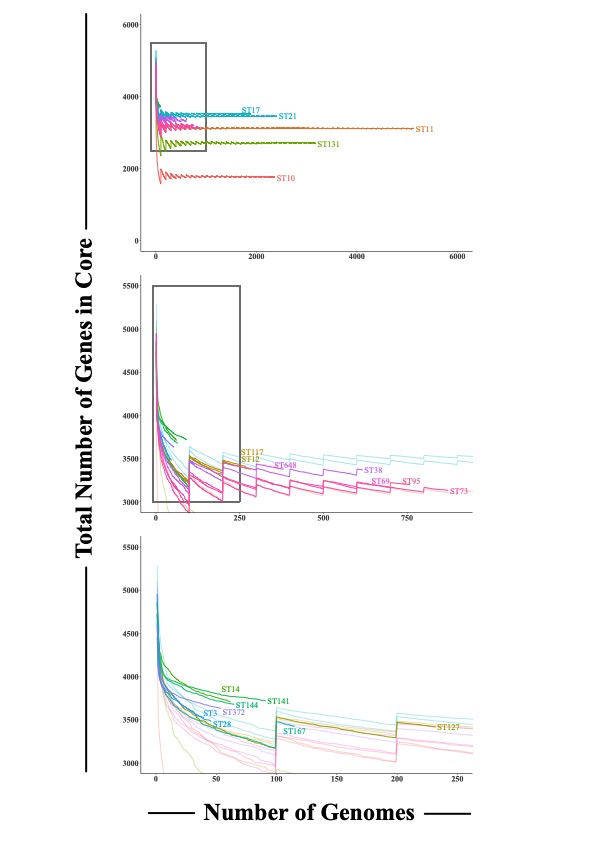

Figure S15: Core genome rarefaction curves for each ST.

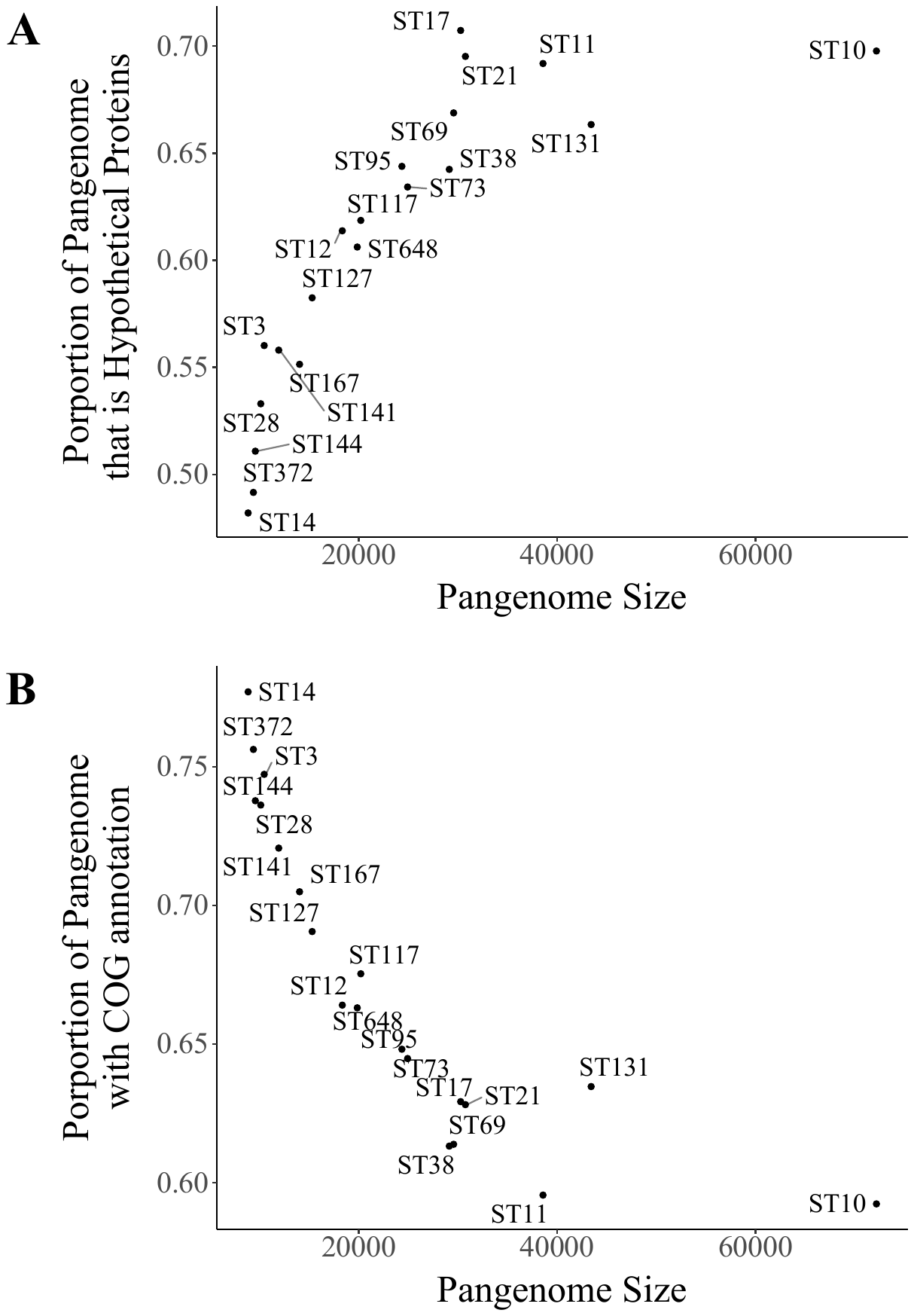

Figure S16: A – Pangenome total size against the proportion of genes annotated as hypothetical proteins for each ST. B- - Pangenome total size against the proportion of the pangenome assigned a COG category for each ST.
